# Evolution of plant cell-type-specific *cis*-regulatory elements

**DOI:** 10.1101/2024.01.08.574753

**Authors:** Haidong Yan, John P. Mendieta, Xuan Zhang, Alexandre P. Marand, Yan Liang, Ziliang Luo, Mark A.A. Minow, Hosung Jang, Xiang Li, Thomas Roulé, Doris Wagner, Xiaoyu Tu, Yonghong Wang, Daiquan Jiang, Silin Zhong, Linkai Huang, Susan R. Wessler, Robert J. Schmitz

## Abstract

*Cis*-regulatory elements (CREs) are critical in regulating gene expression, and yet understanding of CRE evolution remains challenging. Here, we constructed a comprehensive single-cell atlas of chromatin accessibility in *Oryza sativa*, integrating data from 103,911 nuclei representing 126 discrete cell states across nine distinct organs. We used comparative genomics to compare cell-type resolved chromatin accessibility between *O. sativa* and 57,552 nuclei from four additional grass species (*Zea mays, Sorghum bicolor, Panicum miliaceum*, and *Urochloa fusca*). Accessible chromatin regions (ACRs) had different levels of conservation depending on the degree of cell-type specificity. We found a complex relationship between ACRs with conserved noncoding sequences, cell-type specificity, conservation, and tissue-specific switching. Additionally, we found that epidermal ACRs were less conserved compared to other cell types, potentially indicating that more rapid regulatory evolution has occurred in the L1-derived epidermal layer of these species. Finally, we identified and characterized a conserved subset of ACRs that overlapped the repressive histone modification H3K27me3, implicating them as potentially silencer-like CREs maintained by evolution. Collectively, this comparative genomics approach highlights the dynamics of plant cell-type-specific CRE evolution.

## Main

*Cis*-regulatory elements (CREs) function as pivotal hubs, facilitating the binding of transcription factors (TFs) and recruitment of chromatin-modifying enzymes, thereby fine-tuning gene expression in a spatiotemporal-specific manner^1^. CREs play important roles in developmental and environmental processes, and their functional divergence frequently drives evolutionary change^2,3^. Prior studies highlighted the dynamic nature of CREs throughout evolution and their involvement in regulating gene expression via distinct chromatin pathways^4-9^. Across diverse cell types, gene expression is intricately regulated by multiple distinct CREs, each exerting control within specific cell, tissue type, particular developmental stage, or environmental cue^10-12^. In plants, environmental sensing and adaptation relies heavily upon epidermal cells^13^. For example, grass epidermal bulliform cells change their turgor pressure to roll the leaf to slow water loss under stressful conditions, with the TF, ZINC FINGER HOMEODOMAIN 1 (ZHD1), modulating leaf rolling by influencing rice (*Oryza sativa*) bulliform cell development^14,15^. Several studies have identified CREs functioning in a cell-type-specific manner within diverse plant species^16-23^. Despite these findings, our understanding of CREs exhibiting evolutionarily conserved or divergent cell-type-specific activities remains limited.

Through single-cell assay for transposase accessible chromatin sequencing (scATAC-seq), we constructed an expansive single-cell reference atlas (103,911 nuclei) of accessible chromatin regions (ACRs) within rice. We then leveraged these data in tandem with four additional scATAC-seq leaf datasets from diverse grasses [*Zea mays* (16,060 nuclei), *Sorghum bicolor* (15,301 nuclei), *Panicum miliaceum* (7,081 nuclei), and *Urochloa fusca* (19,110 nuclei)]^24^ allowing us to compare ACRs across species and cell types. We quantified the proportion of ACRs that were conserved in these monocots and found high rates of cell-type-specific ACR turnover, particularly in epidermal cells. This indicates that the ACRs associated with specific cell types are rapidly evolving. Finally, we used both conserved non-coding sequences (CNS) and H3K27me3 to find a series of conserved ACRs and the candidate CREs within them that are potentially important for recruitment of Polycomb-mediated gene silencing.

### Construction of an ACR atlas at single-cell resolution in rice

To create a cell-type-resolved ACR rice atlas, we conducted scATAC-seq across a spectrum of nine organs in duplicate (Fig. 1a and b). Data quality metrics, such as correlation between biological replicates, transcription start site enrichment, fraction of reads in peaks, fragment size distribution, and organelle content, revealed excellent data quality (Supplementary Fig. 1 and 2). Following strict quality control filtering, we identified 103,911 high-quality nuclei, with an average of 41,701 unique Tn5 integrations per nucleus. Based on a nine-step annotation strategy, which included RNA *in situ* and spatial-omic (slide-seq) validation of cell-type specificity, we identified a total of 126 cell states, encompassing 59 main cell types across various developmental stages from all the organs sampled (Fig. 1b; Extended Data Fig. 1a; Supplementary Note 1; Supplementary Fig. 3-13; Supplementary Tables 1-7).

**Fig. 1.**
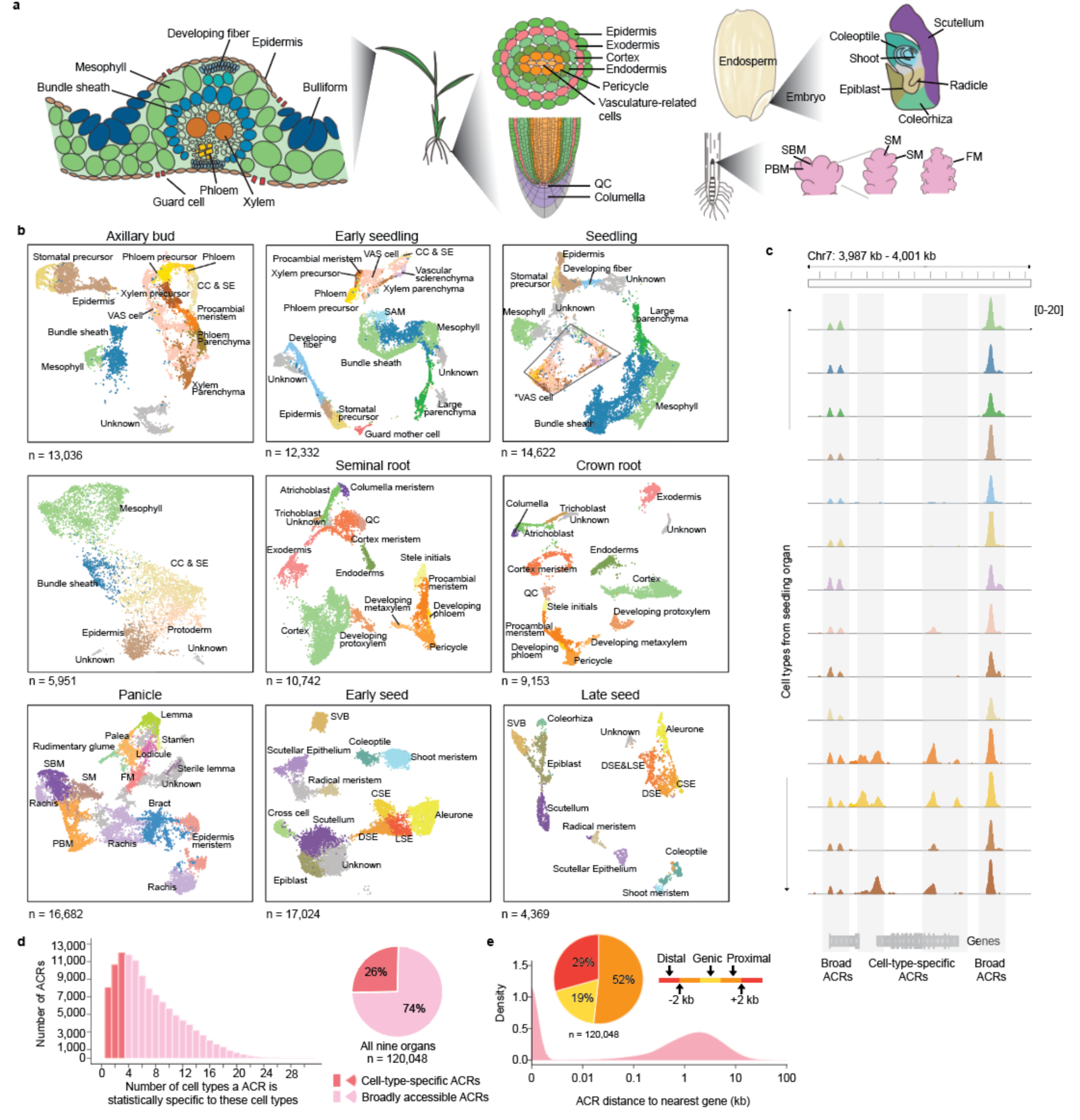
Identifying cell types and characterizing ACRs in rice using scATAC-seq data. **a,** Overview of cell types in leaf, root, seed, and panicle organs. QC: Quiescent Center. SBM: Secondary Branch Meristem. PBM: Primary Branch Meristem. SM: Spikelet Meristem. FM: Floral Meristem. **b,** UMAP projection of nuclei, distinguished by assigned cell-type labels in axillary bud, early seedling (7 days after sowing), seedling (14 days after sowing), leaf (V4 stage; four leaves with visible leaf collars), seminal root, crown root, panicle, early seed development (6 DAP; days after pollination), and late seed development (10 DAP). SAM: Shoot Apical Meristem. VAS cells: vasculature-related cells. *VAS cells: vasculature-related cells, which were further distinguished as procambial meristem, developing phloem/xylem, developing phloem/xylem precursor, vascular parenchyma/sclerenchyma, xylem parenchyma, and companion cell & sieve elements in Supplementary Fig. 6. CC & SE: Companion cell & sieve elements. CSE: Central Starchy Endosperm. DSE: Dorsal Starchy Endosperm. LSE: Lateral Starchy Endosperm. SVB: Scutellar Vascular Bundle. **c,** Evaluation of proportions of ACRs that are cell-type specific versus broad. **d,** A screenshot illustrates the examples of cell-type-specific and broad ACRs. **e,** Accessible chromatin regions (ACRs) show a bimodal distribution of distance to the nearest gene. The ACRs were categorized into three major groups based on their locations to the nearest gene: genic ACRs (overlapping a gene), proximal ACRs (located within 2 kb of genes), and distal ACRs (situated more than 2 kb away from genes).

By analyzing cell-type-aggregated chromatin accessibility profiles, we identified a total of 120,048 ACRs (Extended Data Fig. 1b and c). Among these ACRs, 30,796 were categorized as ‘cell-type-specific ACRs’, exhibiting cell-type-specific entropy signals of accessible chromatin in less than 5% (3/59) of the main cell types, whereas approximately 89,252 were classified as ‘broad ACRs’ with chromatin accessibility in more than 5% of the cell types (Fig. 1c and d; Extended Data Fig. 1d and e). When analyzing ACR proximity to genomic features in the rice genome, about half of the ACRs were gene proximal (52%; located within 2 kb of genes; Fig. 1e). These proximal ACRs had higher but less variable chromatin accessibility than genic and distal ACRs (Extended Data Fig. 1f). In contrast, about 19% of the ACRs overlapped genes, mostly in introns, and the remaining 29% were categorized as distal (Fig. 1e; situated more than 2 kb away from genes). The greater chromatin accessibility variance in non-proximal ACRs suggests these regions may act in select cellular contexts. To further investigate the association of distal ACRs with gene activity, we examined the interactions between distal cell-type-specific ACRs and genes using leaf bulk Hi-C data^25^. Among the 3,513 distal cell-type-specific ACRs in leaf tissue, most (81.7%) were embedded within chromatin loops. More than one-third (37.7%) of these ACRs interacted with promoters of cell-type-specific accessible genes, and 11.2% (392) had both the ACRs and their interacting genes associated with the same cell type (Extended Data Fig. 1g; Supplementary Table 8). As bulk Hi-C poorly measures rare cell types, we expect this number to be a conservative count of the number of cell-type-specific ACRs associated with cell-type-specific gene activity.

### The atlas uncovers key TFs, their motifs, and ACRs during rice development

To demonstrate the utility of this new resource, we associated the atlas ACRs with a set of noncoding quantitative trait nucleotides (QTNs; Fig. 2a). Notably, we observed an enrichment of agriculturally relevant QTNs^26^, within ACRs (Fig. 2b), some of which were cell-type-specific ACRs (Extended Data Fig. 2a and b; Supplementary Table 9). For instance, a QTN was within an endosperm-specific ACR located at ∼1 kb upstream of *GLUTELIN TYPE-A2 PRECURSOR* (*OsGluA2*) (Fig. 2c), which is associated with increased seed protein content^27^. Exploring the endosperm epigenome more, we observed an endosperm-specific reduction in cytosine methylation at endosperm-specific ACRs, including an ACR linked to the DNA demethylase *OsROS1* (Extended Data Fig. 2c)^28^. We found that 5,159 ACRs had lower DNA methylation in the endosperm compared to early seedling (Fig. 2d), and these ACRs were enriched for several TF motif families such as MADS box factors, TEOSINTE BRANCHED1/CYCLOIDEA/PROLIFERATING CELL FACTOR (TCP), Basic Leucine Zipper (bZIP), and BARLEY B RECOMBINANT/BASIC PENTACYSTEINE (BBR/BPC), compared to constitutively unmethylated ACRs (Fig. 2e). Therefore, established endosperm DNA demethylation^29^, coincides with endosperm-specific ACRs.

**Fig. 2.**
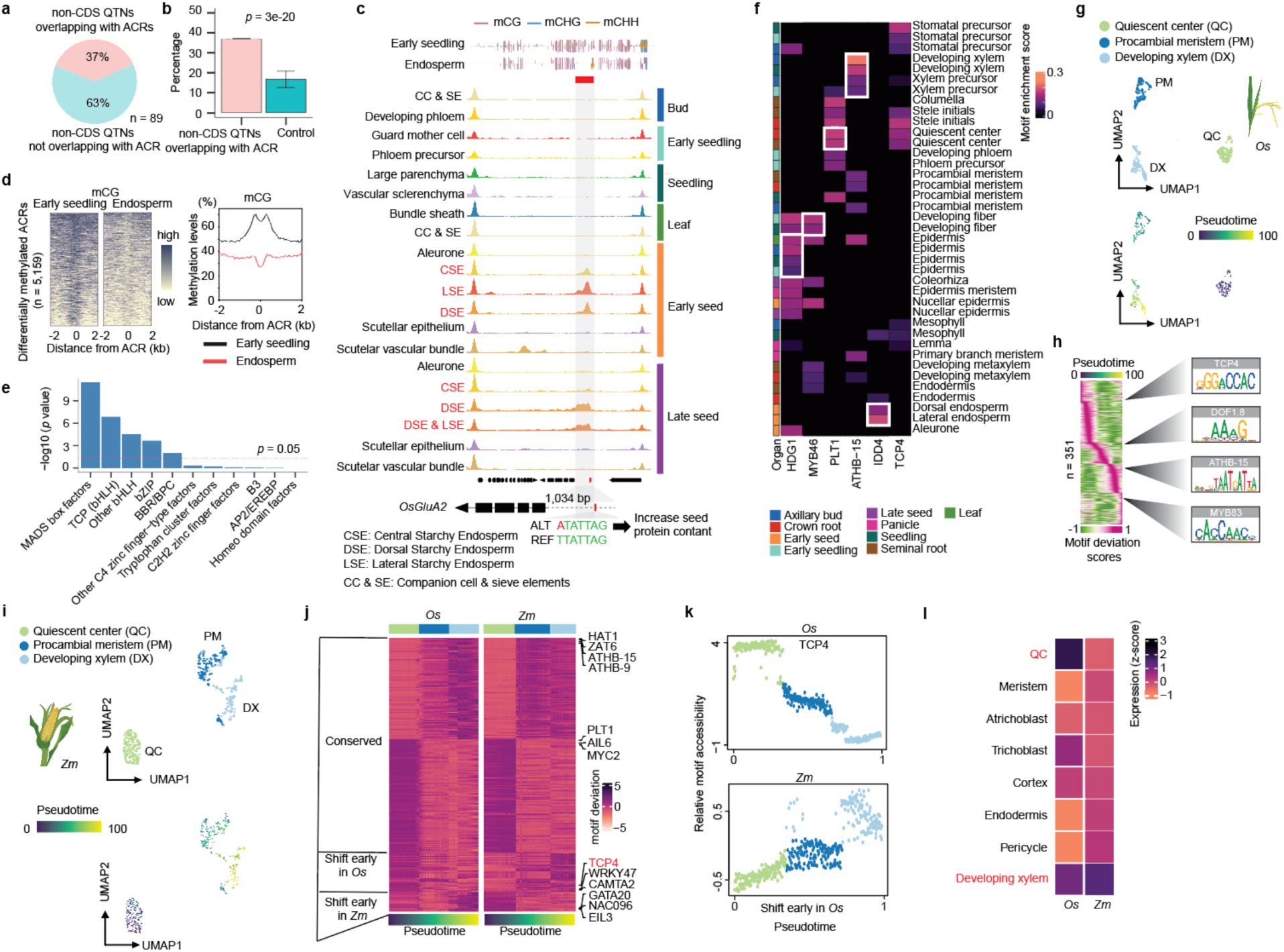
Characterization of TF motifs essential for specific cell types using rice ACR atlas. **a,** The ratio of non-CDS QTNs overlapping with ACRs to all non-CDS QTNs. **b,** A bar plot showing a percentage of non-CDS QTNs overlapping with ACRs. The significance test was done by using the binomial test (alternative = ‘two.sided’; See Methods: Construction of control sets for enrichment tests). **c,** Analysis of cell-type-aggregate chromatin accessibility across 12 seed cell types and eight non-seed-related cell types, showing signatures of a QTN within an endosperm-specific ACR situated at the promoter region of *OsGluA2*. An endosperm-specific reduction of cytosine methylation was identified over the endosperm-specific ACR. **d,** A total of 5,159 differentially methylated ACRs were identified, showing hypermethylation in early seedling tissue but hypomethylation within endosperm tissue. **e,** Enrichment of TF families based on their motifs within the 5,159 ACRs. The upper plot shows the TF motif count within the TF family indicated on the x-axis of the bottom plot. The *p* value was computed using a hypergeometric test (alternative = ‘two.sided’). **f,** Deviations of motifs displaying enrichment in specific cell types (white frames) where their cognate TFs are known to be accumulated. **g,** UMAP visualizations depict the cell progression of root developing xylem in *O. sativa*. **h,** Relative motif deviations for 351 TF motifs (left). Four motifs enriched along the trajectory gradient are shown on the right. **i,** UMAP visualizations depict the cell progression of root developing xylem in *Z. mays*. **j,** The heatmap displays motif deviation scores ordered by pseudotime (x-axis). ‘Conserved’ indicates motifs that showed consistent patterns of motif deviation score changes along the pseudotime between *O. sativa* and *Z. mays*. ‘Shift early’ indicates higher motif deviation scores observed at the beginning of pseudotime. **k,** Comparison of relative motif accessibility of TCP4 along the pseudotime between *O. sativa* and *Z. mays*. **l,** Expression profiles of the *TCP4* gene using scRNA-seq datasets from *O. sativa* and *Z. mays*.

We further investigate the TF motifs that exhibited enrichment within the ACRs of specific cell types (Supplementary Table 10). We observed consistency between known TF activity and cognate motif enrichment (Fig. 2f). For example, ARABIDOPSIS THALIANA HOMEOBOX PROTEIN 15 (ATHB-15) is associated with xylem differentiation^30^ and we found a significant enrichment of the ATHB-15 motif in seedling developing xylem or xylem precursor cells. Other TFs, such as HOMEODOMAIN GLABROUS 1 (HDG1)^31^, MYELOBLASTOSIS 46 (MYB46)^32^, PLETHORA1 (PLT1)^33^, INDETERMINATE DOMAIN 4 (IDD4)^34^ are known to accumulate in the epidermis, developing fiber, quiescent center (QC), and endosperm, respectively, and their motifs were enriched in these cell types (Fig. 2f). Furthermore, the motif analysis unveiled potential novel roles for certain cell types. For instance, AtTCP4 regulates lignin and cellulose deposition and binds the promoter of *VASCULAR-RELATED NAC DOMAIN 7* (*VND7*), a pivotal xylem development gene^35^. However, in *O. sativa*, the TCP4 motif was QC enriched more so than in the developing root xylem, alluding to unknown QC roles. In sum, our TF motif enrichment sheds light on both known and novel regulatory mechanisms underlying cell differentiation and function.

To examine the ACR dynamics during cell fate progression, we organized nuclei along pseudotime trajectories representing 14 developmental continuums (Fig. 2g; Extended Data Fig. 2d and e; Supplementary Fig. 14; Supplementary Table 11 and 12), focusing on root developing xylem (RDX), we identified 16,673 ACRs, 323 of 2,409 TFs, and 351 of 540 TF motifs showing differential chromatin accessibility along the xylem trajectory (Supplementary Table 13). Early in the xylem trajectory, the TCP4 motif was notably enriched (Fig. 2h). To determine if TCP4 enrichment is also present in *Z. mays*, we aligned the RDX motifs of both species using a dynamic time-warping algorithm (Extended Data Fig. 2f), which identified 62 motifs with species-differential *cis*-regulatory dynamics during RDX developments (Extended Data Fig. 2f; Supplementary Table 14). OsTCP4 decreased in motif accessibility during xylem development, whereas ZmTCP4 increased along RDX trajectory (Fig. 2j and k). This mirrors the single-cell RNA sequencing (scRNA-seq) expression patterns of *TCP4* during RDX development^36^ (Fig. 2l). This unveiling of opposing developmental TCP4 motif accessibility gradients in *O. sativa* and *Z. mays,* exemplifies how our atlas can merge with existing, and future data, to drive discovery surrounding monocot development.

In sum, the *O. sativa* atlas provides a comprehensive ACR resource, capturing known agronomic QTNs and bringing novel insights to seed and xylem development. Beyond the discoveries outlined here, this atlas represents a potent reference for the rice research community to answer diverse questions about cell type specific processes.

### The landscape of cell-type-specific ACRs across grass species

We were interested in leveraging the *O. sativa* atlas to understand how ACRs change during grass evolution. The *O*. *sativa* ACRs were overlapped with syntenic regions defined by their relationship to four different grass species *Z. mays*, *S. bicolor*, *P. miliaceum*, and *U. fusca* that have single-cell ATAC sequencing using combinatorial indexing (sciATAC-seq) data from leaves^24^. The analysis revealed that 34% (40,477) of the *O. sativa* ACRs were within 8,199 syntenic regions (∼86 Mb of the *O. sativa* genome) shared with at least one of the four examined grass species (Extended Data Fig. 3a; Supplementary Fig. 15). In contrast, the majority of ACRs (66%; 79,571) were in non-syntenic regions. Notably, the ACRs found in non-syntenic regions were significantly enriched for cell-type specificity ACRs (*p =* 6e-248 to 0.0031; Fisher’s exact test), for both proximal and distal ACRs (Extended Data Fig. 3b and c). This reveals that most of the *O. sativa* ACRs in the grass species examined occurred in non-syntenic regions.

To determine to what degree ACR number, genomic position and cell-type specificity differs amongst grasses, we compared the composition and distribution of leaf ACRs across the five species (Fig. 3a). Using previous cell-type annotations^24^, we calculated the proportion of both broad and cell-type-specific ACRs across all species. We categorized cell-type-specific ACRs as those only accessible in 1-2 leaf cell-types (Extended Data Fig. 3d; Supplementary Table 15). An average of ∼53,000 ACRs were identified across the five species, with 15-35% of the ACRs classified as cell-type specific (Fig. 3b; Supplementary Fig. 16 and 17). Broad and cell-type-specific ACRs were equivalent in their distributions around promoters, distal, and genic regions (Fig. 3c; Extended Data Fig. 3e-f).

**Fig. 3.**
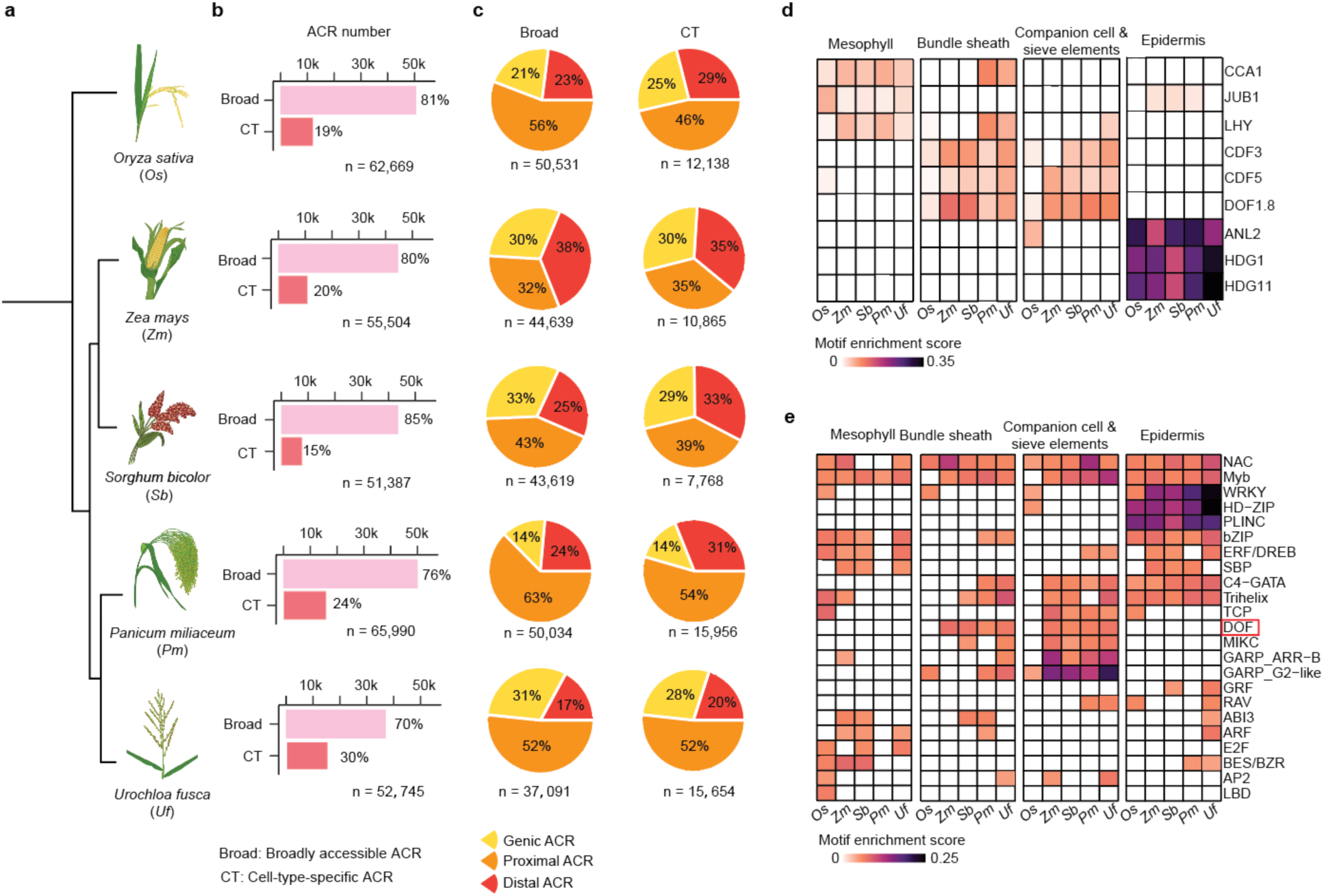
Position and motif enrichment of cell-type-specific ACRs across species. **a,** A phylogenetic tree illustrates five species under examination. **b,** The count of broad and cell-type-specific ACRs. **c,** Broad and cell-type-specific ACRs were classified into three main groups based on their proximity to the nearest gene: genic ACRs (overlapping a gene), proximal ACRs (located within 2 kb of genes), and distal ACRs (situated more than 2 kb away from genes). *O. sativa*, *P. miliaceum*, and *U. fusca* showed a higher percentage of proximal ACRs, but a lower percentage of distal ACRs compared to *Z. mays* and *S. bicolor*, likely reflecting differences in intergenic space and overall genome sizes. **d**, A heatmap illustrates nine TF motif enrichments, consistent with the known TF dynamics among cell types [CIRCADIAN CLOCK ASSOCIATED 1 (CCA1), LATE ELONGATED HYPOCOTYL (LHY) and JUNGBRUNNEN1 (JUB1): mesophyll^43^; CYCLING DOF FACTOR 3 (CDF3) and CDF5: companion cells^44^; DOF1.8: vascular-related cells^45^; ANTHOCYANINLESS2 (ANL2), HOMEODOMAIN GLABROUS 1 (HDG1), and HDG11: epidermis^31,46,47^]. **e,** A heatmap illustrates collapsed TF motif enrichment patterns into super motif families across various species for each cell type. The motif enrichment score cutoff was set to 0.05. The score for each super TF motif family was calculated by averaging the enrichment scores of all the TF motif members within that super family. The DOF TF motif family was highlighted by a red frame. To mitigate the impact of substantial variations in cell numbers across species or cell types, we standardized (down-sampled) the cell counts by randomly selecting 412 cells per cell type per species. This count represents the lowest observed cell count for a given cell type across all species (See Methods: Linear-model based motif enrichment analysis).

Prior hypotheses suggested that large scale regulatory rewiring could play a key role in cell-type environmental adaptation^8,37^. To explore instances where divergent TF activity occurred in the same cell types, we associated TF gene body chromatin accessibility with their cognate TF motifs across different species and cell types. Approximately 64% to 76% of the TFs (211 to 232) examined exhibited a positive correlation between the local chromatin accessibility of their gene body and global enrichment of their cognate binding motifs within ACRs (motif deviation) across all leaf cell types and all species (Extended Data Fig. 4a). The use of TF chromatin accessibility was supported by an analysis of TF expression and motif deviation in both seedling (Supplementary Fig. 7) and root data (Zhang et al. 2021), which uncovered a similar positive relationship across cells (Extended Data Fig. 4b). These results suggest a positive relationship between TF gene-body chromatin accessibility/expression and TF activity in the same cell type.

Moreover, the genomic sequences from all ACRs discovered in all species and cell types exhibited enrichment of TF motifs compared to a control set of sequences (Extended Data Fig. 4c). Furthermore, TF motif enrichment analysis revealed known TF-cell-type specificities (Fig. 3d). For example, the HDG1 TF is critical for epidermis and cuticle development^31^ and its motif was enriched in epidermis cells in all five species (Extended Data Fig. 4d). We also observed motif enrichments of WRKY, HOMEODOMAIN-LEUCINE ZIPPER (HD-ZIP), and PLANT ZINC FINGER (PLINC) in epidermal cells across all five species examined (Fig. 3e; Extended Data Fig. 4e; Supplementary Fig. 18; Supplementary Table 16). This result indicates that these TFs play a conserved role in the development of the grass epidermis. Phloem companion, sieve element cell, and bundle sheath cell TFs exhibited similar enrichments across the species (Fig. 3e). However, species-specific motif patterns were also observed, with *O. sativa* being the most different. For example, the DNA-BINDING ONE ZINC FINGER (DOF) TF family motif exhibited higher enrichment scores in four C_4_ photosynthesizing species (*Z. mays*, *S. bicolor*, *P. miliaceum*, and *U. fusca*), as opposed to C_3_ photosynthesizing *O. sativa* (Fig. 3e; Extended Data Fig. 4f and g). The DOF TF family is involved in *Arabidopsis thaliana* vasculature development^38^, and is important in the transition from C_3_ to C_4_ photosynthesis^23,39-42^. The enrichment of DOF motifs in *O. sativa* species-specific motifs is, therefore, an expected biological signal, indicating robust species-specific motif detection and high-quality single-cell data.

Taken together, our findings demonstrate the power of scATAC-seq data in a comparative framework to explore regulatory evolution, both based on the relationship of ACRs to TF motifs, as well as the relationship between TFs and their corresponding motifs.

### Species-specific evolution of cell-type-specific ACRs

To understand how cell-type-specific and broad ACRs changed over evolution, we examined ACRs within syntenic regions among the studied species. To compare ACRs, we devised a synteny-based BLASTN pipeline that allowed us to compare sequences directly (See Methods: Identification of syntenic regions; Fig. 4a; Extended Data Fig. 5a; Supplementary Table 17). Using *O. sativa* ACRs as a reference, we identified three classes of cross-species ACR conservation: 1) ACRs with matching sequences that are accessible in both species (shared ACRs), 2) ACRs with matching sequences, but are only accessible in one species (variable ACRs), and 3) ACRs where the sequence is exclusive to a single species (species-specific ACRs; Fig. 4b; Extended Data Fig. 3d). The shared ACR BLASTN hits were often small syntenic sequences, highlighting the large divergence of grass ACRs sequences. However, the majority (92-94%) of these shared BLASTN sequences encoded known TF motifs (Supplementary Fig. 18), indicating that shared ACRs are conserved regulatory regions. In contrast, variable ACRs represent a blend of conserved and divergent regulatory elements, and species-specific ACRs likely indicate novel regulatory loci. We found that, on average, shared ACRs were enriched (*p* = 9e-50 to 4e-17; Fisher’s exact test) for broad ACRs, whereas the variable (*p* = 2e-23 to 2e-04; Fisher’s exact test) and species-specific (*p* = 2e-06 to 1e-02; Fisher’s exact test) classes were enriched for cell-type specificity (Fig. 4c; Extended Data Fig. 5b and c). Moreover, we observed that the genomic distribution of shared ACRs were biased towards proximal ACRs (Extended Data Fig. 5d). This contrasts with cell-type-specific ACRs, which are overrepresented in the species-specific class (Fig. 4c). The cell-type-specific ACRs within the species-specific class were more likely to reside in distal genomic regions compared to the ACRs within the shared and variable classes (Extended Data Fig. 5e).

**Fig. 4.**
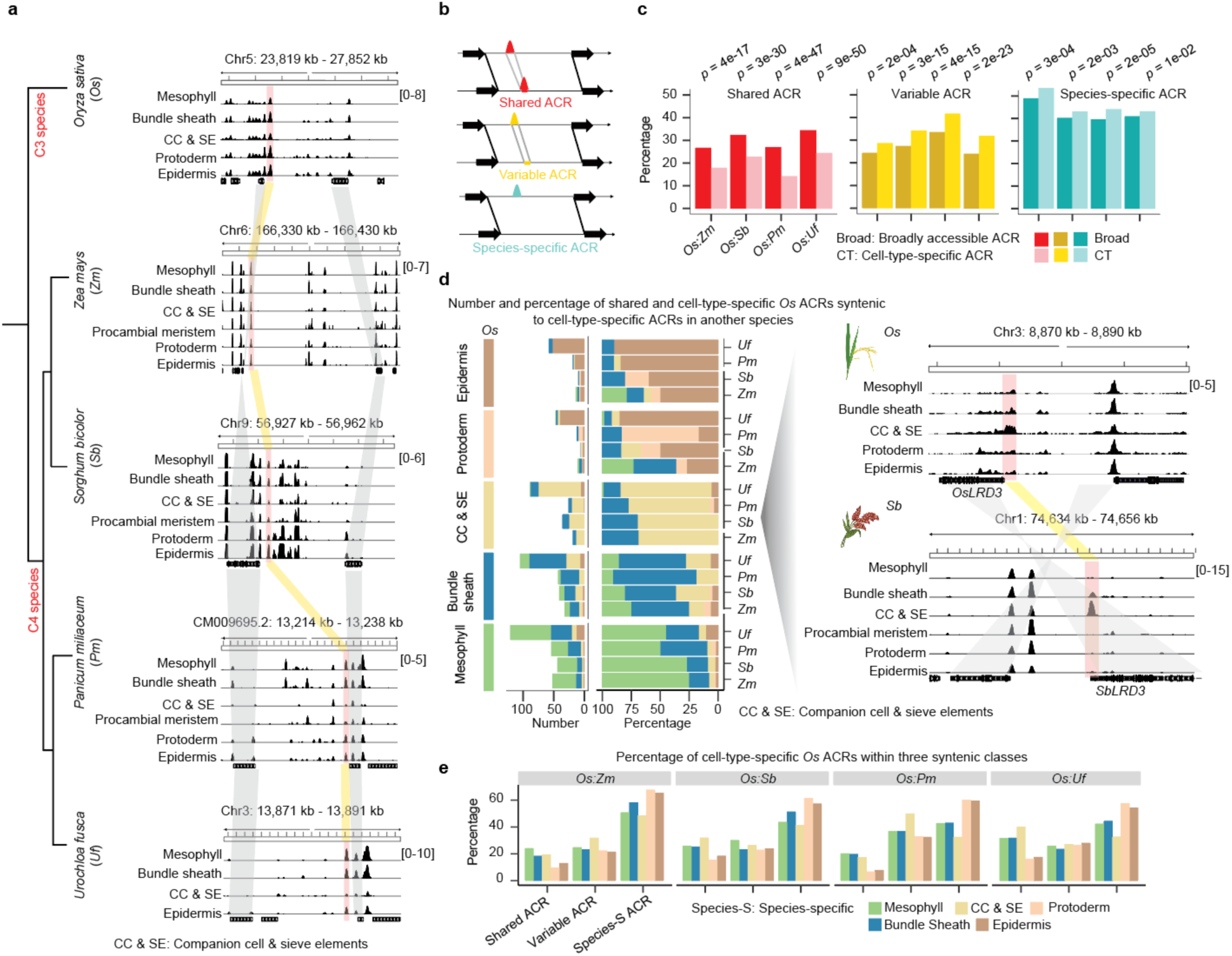
Cell-type-specific ACRs are frequently species-specific. **a,** A screenshot illustrating syntenic regions capturing shared ACRs across five species. The red bars denote syntenic ACRs within regions flanked by corresponding syntenic gene pairs, while the gray color highlights these syntenic gene pairs. **b,** Three classes depicting variations in ACR conservation between two species. ‘Shared ACRs’: ACRs with matching sequences that are accessible in both species; ‘Variable ACRs’: ACRs with matching sequences but are only accessible in one species; ‘Species-specific ACRs’: ACRs where the sequence is exclusive to a single species. **c,** The percentage of broad and cell-type-specific ACRs underlying three classes shown in panel **b**. The significance test was done by using the Fisher’s exact test (alternative = ‘two.sided’). **d, Left**, the number and percentage of *O. sativa* shared ACRs that retain or change cell-type specificity amongst the other four species. **Right**, a screenshot of a *O. sativa* phloem-specific ACR that retains phloem specificity in *S. bicolor*. This ACR is situated at the promoter region of *LRD3* which is specifically expressed in companion cell and phloem sieve elements (Supplementary Table 2). The gray shaded region highlights the syntenic gene pair. **e,** The percentage of cell-type-specific ACRs identified across all cell types within each species pair split into three classes shown in panel **b**. The percentage for each cell type within the three classes collectively sum to 100%.

We further investigated whether the cell-type-specific ACRs were conserved in their cell-type specificity by evolution. Pairwise comparison between *O. sativa* and the other grasses revealed that between 0.7% (137/19,941) to 1.8% (420/22,881) of the syntenic ACRs were shared ACRs retaining the same cell-type specificity in both species (Extended Data Fig. 5b and f). Of these few shared cell-type specific ACRs, the majority (62%-69%), were accessible in the identical cell-type in both *O. sativa* and the corresponding species (Fig. 4d; Extended Data Fig. 5f). For example, the promoter ACR associated with *LATERAL ROOT DEVELOPMENT 3* (*LRD3*), a gene critical in companion cell and sieve element development^48^, showed sequence conservation between *O. sativa* and *S. bicolor* (Fig. 4d). Interestingly, ACRs which were mesophyll specific in *O. sativa* changed their cell-type specificity to bundle sheath 17%-41% of the time, while bundle sheath ACRs changed to mesophyll 9%-25% of the time (Fig. 4d; Extended Data Fig. 5g). This result is likely due to the functional divergence associated with the shift from C_3_ (*O. sativa*) to C_4_ (all other species sampled) photosynthesis^23,24^. Within variable ACRs, leaf companion cell and sieve elements cells were consistently over-represented (Fig. 4e), potentially due to sampling leaves at different stages of the source-sink transition. Of all the classes of cross-species ACR conservation, species-specific ACRs were the most predominant in every cell type (Fig. 4e). These findings suggest a dynamic and rapid evolution of cell-type-specific ACRs within the examined species. Notably, ACRs in L1-derived layer (epidermis and protoderm) exhibited the highest proportion of species-specific ACRs (Fig. 4e; Extended Data Fig. 6a-c). The high divergence of ACRs in L1-derived cells was also observed in the pairwise comparison between *O. sativa* and *Z. mays* seedling (Extended Data Fig. 6d and e), as well as root cells (Extended Data Fig. 6f and g), suggesting that grass ACRs in L1-derived cells are surprisingly divergent compared to internal cell types. Further, we examined *O. sativa* (Supplementary Fig. 7; Supplementary Table 18) and *Z. mays* single-nucleus RNA sequencing (snRNA-seq) data^20^, to investigate whether the L1-derived cell types exhibited the most divergent transcriptomes. We found that, among the six examined cell types, protoderm showed the lowest similarity in the gene expression levels between the two species (Extended Data Fig. 6h), which suggests ACR divergence in L1-derived cells likely drives transcriptional change.

We further investigated the divergence of ACRs in L1-derived cells between more closely related species pairs: *Z. mays* and *S. bicolor*; *P. miliaceum* and *U. fusca*. More closely related species shared more ACRs across all cell types, containing fewer species-specific ACRs (Extended Data Fig. 6i). Unlike when compared to the *O. sativa* reference (Fig. 4e; Extended Data Fig. 6a-c), we did not observe more species-specific ACRs in L1-derived cells compared to other cell-types (Extended Data Fig. 6j). Taken together, we observed that ACRs in L1-derived cells are the most divergent compared to other cell types, but only when comparing grasses over long evolutionary distances.

Returning to *O. sativa* as the reference, we investigated the TF families underpinning the species-specific ACRs in L1-derived cells. Within all species-specific syntenic ACRs, we observed a predominance of TF motifs for the HD-ZIP, SQUAMOSA PROMOTER BINDING PROTEIN (SBP), PLINC families (Extended Data Fig. 7a). Many of these, such as HDG1, ZHD1, ATHB-20, SPL3, SPL4, and SPL5, function in epidermal cell development^15,49-52^. The predominance of these motifs in the species-specific ACR class suggests that although L1 TF motif sequences are well conserved (Fig. 3d and e), the ACRs which contain these motifs are not conserved in grass genomes. Upon comparing TF-motif enrichment in syntenic and non-syntenic ACRs, we observed the presence of these epidermal motif families in both groups (Extended Data Fig. 7b; Supplementary Table 19), indicating their essential roles in both conserved epidermal cell development and rapid gene-regulatory co-option in species-specific sequences. Notably, some TF-motif families, such as WRKY, were more enriched in non-syntenic ACRs in epidermal cells (Extended Data Fig. 7b), further supporting that this family may drive phenotypic innovation in the epidermal layer.

To look for derived species-specific ACRs associated with the altered expression of surrounding gene orthologs in epidermal cells, we integrated snRNA-seq data from *O. sativa* (Supplementary Fig. 7), with a snRNA-seq data from *Z. mays*^20^. We identified 87 orthologous genes, irrespective of synteny, which exhibited higher L1 *O. sativa* expression compared to *Z. mays* (Extended Data Fig. 7c; Supplementary Table 20). A gene ontology enrichment test for these 87 genes revealed eight genes that were significantly enriched in the lipid metabolic process (Extended Data Fig. 7d), possibly related to cuticle metabolism. Among the eight genes, one was orthologous to *A. thaliana GDSL LIPASE GENE* (*LIP1*), which is epidermis specific^53^. We further identified 102 L1 cell-type-specific ACRs from *O. sativa* that were the closest to the 87 orthologous genes and observed 11 TF motifs enriched (*q* = 3e-10 to 5e-04; Binomial test) in these ACRs (Extended Data Fig. 7c). These included TF family motifs known for their roles in epidermal cell development such as ZHD1^15^, HDG11^54^, ZHD5^55^, HDG1^31^, and WRKY25^56^. For example, within the *OsLIP1* intron, we identified two ZHD1 motifs within a species-specific ACR that was specifically accessible in L1-derived cells (Extended Data Fig. 7e). We also flipped this comparison by identifying 166 orthologs with elevated *Z. mays* epidermal expression compared to *O. sativa* (Supplementary Table 20). This revealed 196 L1 cell-type-specific ACRs in *Z. mays*. Within these ACRs, the most enriched (*q* = 0.0129 to 0.0392; Binomial test) TF motif was MYELOBLASTOSIS 17 (MYB17; Extended Data Fig. 7f and g). This R2R3 MYB family TF is associated with epidermal cell development, specifically in the regulation of epidermal projections^57^. Furthermore, we hypothesized that some of these novel motifs could be related to *O. sativa* transposable element (TE) expansion. Notably, we found long terminal repeat retrotransposon (LTR)-associated ACRs from the *Gypsy* family were enriched (*p* = 0.0012 to 0.0415; Fisher’s exact test) in *O. sativa* epidermal cell-type-specific ACRs (Extended Data Fig. 7h). The ZHD1 motif was enriched within these *Gypsy*-associated ACRs (*p* = 0.0011 to 0.0052; Binomial test) (Extended Data Fig. 7i). By linking snRNA-seq to scATAC-seq data, we tied gene-proximal ACR changes to elevated epidermal expression of a small number of conserved orthologs 50 million years derived^58^. These ACR changes are associated with variance in species-specific L1-derived layer development, potentially contributing to species differences in environmental adaptation.

### CNS are enriched in cell-type-specific ACRs

To augment our syntenic ACR BLASTN approach, we intersected our ACRs with published CNS^6,59-61^. Outside of untranslated regions (UTRs), CNS typically encompass transcriptional regulatory sequences undergoing purifying selection, too critical to be lost during evolution^62^. Using the conservatory database (https://conservatorycns.com/dist/pages/conservatory/about.php)^63^, we extracted 53,253 and 284,916 CNS in *O. sativa* and *Z. mays*, respectively, for analysis. Excluding CNS overlapping with UTRs, 30.8% and 21.3% of CNS overlapped with the leaf-derived ACRs in *O. sativa* and *Z. mays*, respectively (Fig. 5a; Extended Data Fig. 8a). Using all ACRs in the *O. sativa* atlas, this ratio increased to 65.0% (Fig. 5a), indicating that a significant portion of these CNS likely function in specific cell types and tissues. One common assumption is that CNS which overlap ACRs (CNS ACRs) retain similar function between species^64-70^. This potential of conserved function makes CNS tempting targets for genome editing, as changes to these sequences can yield morphological variation, some that are important for crop improvement^71,72^. To test assumptions that CNS within ACR classes are ideal targets for genetic modification, we assessed conservation of cell-type specificity associated with CNSs. We observed 39% to 51% of total CNS ACRs within the ‘shared CNS ACR’ class (Extended Data Fig. 8b and c), suggesting these ACRs have conserved cellular contexts between *O. sativa* and other species. We also compared shared ACRs between *O. sativa* and *Z. mays*, both containing CNS. We found that ACRs with identical cell-type specificity had significantly (*p* = 0.04114; Wilcoxon signed rank test) longer alignments and higher (*p* = 3e-05; Wilcoxon signed rank test) TF motif numbers than ACRs with differing cell-type specificity (Extended Data Fig. 8d). Within syntenic regions, CNS ACRs are more cell-type specific than those without (Fig. 5b). This enrichment was consistent for all classes of ACRs we identified, except for species-specific ACRs which were equivalently cell-type specific and broad (Fig. 5b). The enrichment of CNS in cell-type-specific ACRs stresses the importance of rare cell-type function, as the cell-type-specific ACRs are preferentially retained during evolution.

**Fig. 5.**
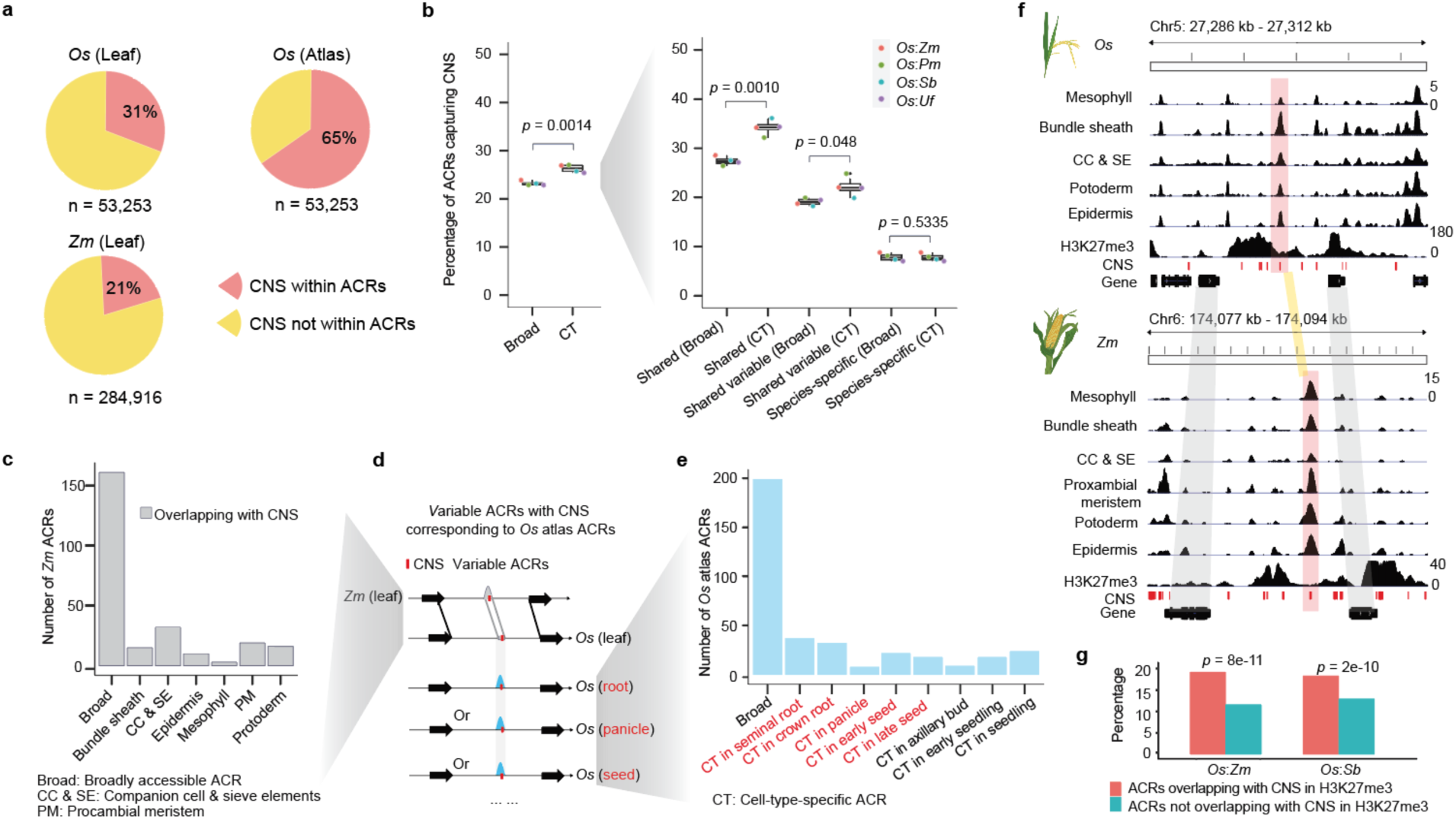
Cell-type-specific ACRs exhibit an enrichment of CNS. **a,** The percentage of CNS overlapping with ACRs. ‘n’ indicates the number of CNS. ‘Atlas’ means the ACRs were from the *O. sativa* atlas in Fig. 1b. **b, Left,** the percentage of broad and cell-type-specific ACRs within syntenic and non-syntenic regions overlapping with the CNS. **Right,** this panel presented similar meaning as the left panel but focusing on three classes within syntenic regions shown in Fig. 4b. Broad: Broadly accessible ACR; CT: Cell-type-specific ACR. Significance testing was performed using the t-test (alternative = ‘two.sided’). **c,** The bar plot showcases the count of *Z. mays* variable ACRs accessible in leaf cell types. **d,** A sketch illustrating whether variable ACRs containing CNS in *Z. mays* capture ACRs derived from the *O. sativa* atlas. **e,** The bar plot represents the count of *O. sativa* atlas ACRs accessible in non-leaf cell types. **f,** An example of a syntenic block containing *O. sativa*-to-*Z. mays* conserved ACRs within a H3K27me3 region. CNS were highlighted using red color. **g,** The percentage of ACRs capturing CNS in and outside of H3K27me3 regions. The percentage for each group within H3K27me3 and not within H3K27me3 regions collectively sum to 100%. Significance testing was performed using Fisher’s exact test (alternative = ‘two.sided’).

However, the majority (49-61%) of all CNS ACRs differed in cell-type specificity between *O. sativa* and *Z. mays* (Extended Data Fig. 8b and c). Specifically, we examined the CNS found in *Z. mays* ACRs that did not have a corresponding leaf ACR in *O. sativa*. Leveraging the *O. sativa* atlas, we identified these sequences had divergent cellular or tissue chromatin accessibilities. 249 (75%) of the *Z. mays* leaf variable CNS ACRs were accessible in non-leaf cell states (Fig. 5c-e; Extended Data Fig. 8e), highlighting the instability of the cellular context in which CNS acts. This suggests frequent co-option of CNS ACRs into different tissues or cell types. Investigating the CNS ACRs that lost leaf cell-type specificity, we observed that these ACRs were accessible in many non-leaf cell types, uniformly distributed amongst the atlas cell annotations (Extended Data Fig. 8f-i). Consistent with our findings that epidermal-specific ACRs tend to have the most species-specific ACRs in syntenic regions (Fig. 4e; Extended Data Fig. 6a-c), L1-derived cells showed a significantly lower ratio of non-syntenic CNS ACRs compared to other cell types (Extended Data Fig. 8j). This lower ratio demonstrates the frequent loss of epidermal CNS, further supporting the rapid evolution of epidermal transcriptional regulation.

Interestingly, we noticed a pattern where some CNS within ACRs also overlapped domains of H3K27me3^6^ (Fig. 5f). H3K27me3 is a histone modification associated with facultative heterochromatin established by the POLYCOMB REPRESSIVE COMPLEX 2 (PRC2)^73-75^. Genes silenced by PRC2 and H3K27me3 are important regulators that are only expressed in narrow developmental stages or under specific environmental stimuli, where they often initiate important transcriptional changes^74,76^. This importance makes the identification of key CREs controlling H3K27me3 silencing especially interesting. Upon closer examination of the ACRs overlapping domains of H3K27me3 to ACRs away from H3K27me3, we observed that H3K27me3 ACRs were significantly enriched for CNS (Fig. 5g). This enrichment supports that some of these CNS underpin conserved, and critical components, of H3K27me3 silencing.

### Candidate silencer CREs are enriched in broad ACRs

To assess the stability and change of H3K27me3 related ACRs across grass lineages, we focused on comparing *O. sativa*, *Z. mays*, and *S. bicolor* using published H3K27me3 ChIP-seq data^6^. We examined ACRs near or within H3K27me3 regions and classified them into two groups: H3K27me3-broad, representing H3K27me3 associated ACRs with chromatin accessibility in many cell types and H3K27me3-cell-type specific, those H3K27me3 associated ACRs with chromatin accessibility in few cell types (Fig. 6a; Extended Data Fig. 3g). The proportion of H3K27me3-broad and H3K27me3-cell-type-specific ACRs was consistent across all species (Extended Data Fig. 9a). H3K27me3-broad ACRs exhibited a depletion of H3K27me3 at the ACR (Fig. 6b), consistent with nucleosome absence in ACRs^77^. In contrast, H3K27me3 depletion was not observed in H3K27me3-cell-type-specific ACRs, with most cells in the bulk ChIP-seq likely containing H3K27me3-modified nucleosomes (Fig. 6b). This is consistent with the H3K27me3-cell-type-specific ACRs potentially acting after the removal of facultative heterochromatin in a specific cell type(s). However, the chromatin accessibility of the H3K27me3-broad ACRs appears to be concurrent with H3K27me3, suggesting these ACRs may regulate H3K27me3 maintenance and removal across most cellular contexts.

**Fig. 6.**
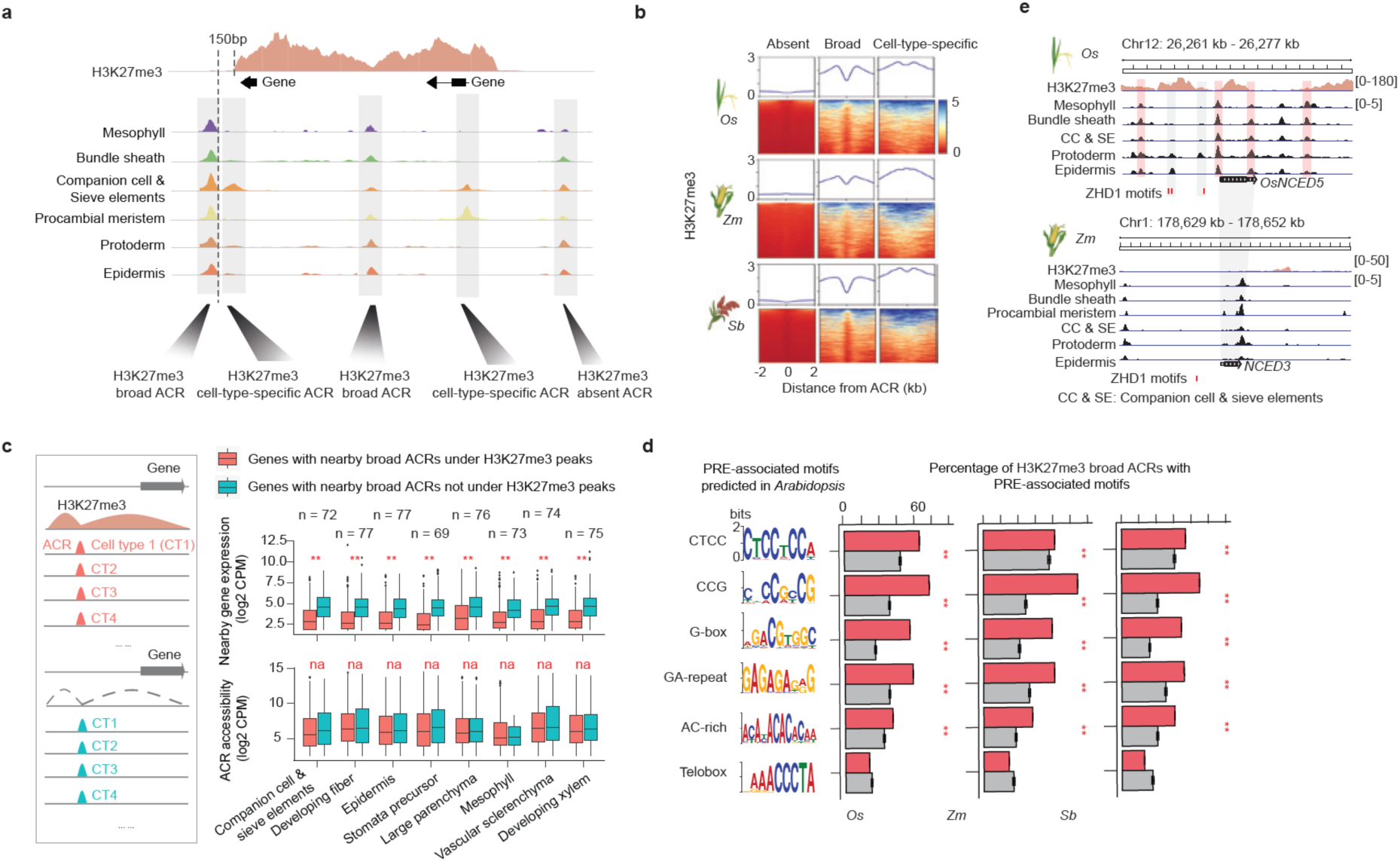
Discovery of candidate silencer CREs across species. **a,** A sketch graph illustrates the classification of ACRs based on their proximity to H3K27me3 peaks. We classified the ACRs into two groups: H3K27me3-associated ACRs (found within or surrounding H3K27me3 peaks) and H3K27me3-absent ACRs. The H3K27me3-associated ACR were further divided into broad ACRs, characterized by chromatin accessibility in at least five cell types, and cell-type-specific ACRs, accessible in less than three out of six examined cell types across all the species. **b,** Alignment of H3K27me3 chromatin attributes at summits of distinct ACR groups. **c,** A comparative analysis of expression levels and chromatin accessibility of genes surrounding broad ACRs under and outside of H3K27me3 peaks. ** indicate *p* value < 0.01, which was performed by the Wilcoxon signed rank test (alternative = ‘two.sided’). The broad ACRs where the H3K27me3 region overlapped >50% of the gene body were positioned within 500 to 5,000 bp upstream of the transcriptional start site of their nearest gene. **d,** Percentage of H3K27me3-broad ACRs in *O. sativa*, *Z. mays*, and *S. bicolor* capturing six known motifs enriched in PREs in *A. thaliana*. ** indicate *p* value < 0.01, which was performed by the Binomial test (alternative = ‘two.sided’; See Methods: Construction of control sets for enrichment tests). **e,** a screenshot of *OsNCED5* accessibility in *O. sativa* and *NCED3* accessibility in *Z. mays* L1-derived cells which contains four H3K27me3-broad ACRs and two *O. sativa* epidermal specific and species-specific ACRs with three ZHD1 motif sites.

To assess the transcriptional state of genes near H3K27me3-broad ACRs, we evaluated snRNA-seq/scRNA-seq from *O. sativa* seedling (Supplementary Fig. 7) and root^78^, and *Z. mays* seedling^20^. The results revealed significantly lower expression (*p* = 3e-34 to 4e-06; Wilcoxon signed rank test) for H3K27me3-broad ACRs associated genes across most cell types (Fig. 6c; Extended Data Fig. 9b and c; Supplementary Table 21). Moreover, 58 bulk RNA-seq libraries from *O. sativa* organs^79^, demonstrated the lower (*p* = 2e-07 to 0.0265; Wilcoxon signed rank test) expression of genes near H3K27me3-broad ACRs than genes near H3K27me3-absent broad ACRs (Extended Data Fig. 9d). To dissect the roles of H3K27me3-broad ACRs, we identified 2,164 H3K27me3-broad ACRs and measured neighboring gene expression in *O. sativa* cells (Supplementary Table 22). About 926 (∼42.8%) of the H3K27me3-broad ACRs were associated with 838 genes that exhibited no expression across any sampled cell type, which was marginally, but significantly (*p* = 1e-10; Fisher’s exact test), higher than the other ACRs associated with the unexpressed genes (Extended Data Fig. 9e; Supplementary Table 22), consistent with these H3K27me3 proximal genes only being expressed under specific conditions. The 1,108 expressed genes associated with H3K27me3-broad ACRs were enriched (*p* < 2e-16; Fisher’s exact test) for cell-type specificity compared to genes without H3K27me3 (Extended Data Fig. 9f). In summary, single-cell expression analysis revealed that the genes linked to H3K27me3-broad ACRs exhibited the hallmarks of facultative gene silencing.

We hypothesized that the H3K27me3-broad ACRs would be enriched for PRC2 silencer elements, as their consistent chromatin accessibility provides an avenue to recruit PRC2 to maintain H3K27me3 throughout development. Supporting the presence of silencer CREs with these ACRs, a known silencer CRE ∼5.3 kb upstream of *FRIZZY PANICLE* was within a H3K27me3-broad ACR^80^ (Extended Data Fig. 10a). To exploit the known Polycomb *A. thaliana* targets, we used scATAC-seq^20^ and H3K27me3^81^ data from *A. thaliana* roots and annotated H3K27me3-broad ACRs. The *A. thaliana* H3K27me3-broad ACRs significantly (*p* < 2e-16; Binomial test) captured 53 of the 170 known Polycomb responsive elements compared to a control class of ACRs, supporting their putative silencer function^74^ (Extended Data Fig. 10b). Furthermore, we implemented a *de novo* motif analysis on the 170 *A. thaliana* elements and identified all reported Polycomb response element (PRE) motifs (CTCC, CCG, G-box, GA-repeat, AC-rich, and Telobox)^74^ (Fig. 6d). Using these motifs and our chromatin accessibility data, we predicted putative binding sites in *O. sativa*, *Z. mays*, and *S. bicolor*, and observed that five motifs were significantly (*p* = 2e-178 to 1e-05; Binomial test) enriched in the H3K27me3-broad ACRs compared to a genomic control (Fig. 6d; Extended Data Fig. 10c). Between 88.0% and 92.7% of H3K27me3-broad ACRs contained at least one PRE motif, with few (0.1-0.2%) ACRs having all six PRE motif types (Extended Data Fig. 10d). Surprisingly, we observed the rates of PRE motif occurrence, and PRE motif counts, were comparable between of H3K27me3-broad ACRs and H3K27me3-absent-broad ACRs (Extended Data Fig. 10c-e). To investigate why the H3K27me3-absent-broad ACRs still contain these motifs, we analyzed ChIP-seq of EMF2b, a crucial PRC2 component^74,82,83^, revealing a significant overlap between EMF2b peaks and the PRE motifs (Extended Data Fig. 10f). Both H3K27me3-broad and H3K27me3-absent-broad ACRs exhibited high EMF2b signals, consistent with the PRE motifs within both ACR groups (Extended Data Fig. 10g). However, H3K27me3-broad ACRs showed higher EMF2b signal in the shoulder (non-peak) areas compared to the H3K27me3-absent-broad ACRs. The EMF2b ChIP-seq suggests PRC2 is recruited to many broadly accessible ACRs, but additional factors are required to activate PRC2 deposition of H3K27me3. In addition, we identified 236 genes and their adjacent H3K27me3-broad ACRs containing PRE motifs with SNPs or Indels in ‘Zhenshan 97’ genotype using ‘Nipponbare’ as a reference in *O. sativa* ^84,85^. A total of 133 of these genes were detected with transcriptional changes between the two genotypes (Supplementary Table 23). In ‘Zhenshan 97’, we observed a significant (*p* = 0.03; Wilcoxon signed rank test) increase in gene expression, accompanied by a decrease in H3K27me3 signal (Extended Data Fig. 10h-j). This suggests that mutations in PRE motifs may preclude PRC2 targeting, potentially associating with higher expression levels of nearby genes within the examined two genotypes. Beyond PRE motifs, we observed significant enrichment (*p* = 1e-16 to 0.0376; Hypergeometric test) of motifs from four TF families in H3K27me3-broad ACRs: APETALA2-like (AP2), basic Helix-Loop-Helix (bHLH), Basic Leucine Zipper (bZIP), and C2H2 zinc-finger (ZnF) (Extended Data Fig. 10k; Supplementary Table 24). AP2 and C2H2 are known to recruit PRC2^74^, and our motif enrichment supports all these TF families potentially regulating H3K27me3 deposition and facultative heterochromatin formation.

To investigate the relationship between these candidate silencers and species divergence, we mirrored our previous syntenic ACR analysis by classifying *O. sativa* H3K27me3-broad ACRs into shared, variable or species specific groups (Extended Data Fig. 11a-c; Supplementary Table 25). Between 54% and 61% of the H3K27me3-broad ACRs were present in the species-specific class, with the H3K27me3-broad ACRs enriched for species-specificity compared to H3K27me3-absent-broad ACRs (Extended Data Fig. 11d). Since these H3K27me3-broad ACRs exhibit hallmarks of PRC2 recruitment, we suspect that altered silencer CREs use context to drive species-specific developmental and environmental responses.

Since we identified L1-derived cells as being enriched in species-specific ACRs, we sought to examine the changes in H3K27me3 regulation within this tissue. We examined our previously identified 87 *O. sativa*-to-*Z. mays* orthologs to see if these genes contained H3K27me3. We observed 18 of these 87 genes were close to H3K27me3-broad ACRs (Supplementary Table 26). For example, we identified four H3K27me3-broad ACRs, and three ZHD1 motifs linking bulliform cell development^14,15^, within two species-specific ACRs surrounding *OsNCED5* that were specifically accessible in L1-derived cells (Fig. 6e). OsNCED5 TF is known to regulate tolerance to water stress and regulate leaf senescence in *O. sativa*^86^. These results highlight that H3K27me3 mediated silencing may play a critical role in divergent regulation in L1-derived cells.

## Discussion

Our comparison of *O. sativa* with four other grasses revealed patterns in the evolutionary dynamics of *O. sativa* ACRs within syntenic and non-syntenic regions and discovered that the grass L1-derived cells exhibit elevated rates of transcriptional regulatory divergence, as well as changes in *cis* regulatory architecture as compared to other cell types over large evolutionary distance (Fig. 7a). However, we revealed a contrasting dichotomy; the epidermal TF motifs were the most cell-type specific of all those studied, yet their cognate ACRs exhibited the strongest target divergence amongst the measured species. This duality highlights tandem conservation of core epidermal motifs, and the rapid co-option of novel regulatory regions into these existing regulatory frameworks. This rapid regulatory evolution might relate to the dynamic environmental pressures the epidermis has evolved to withstand, and may relate to the higher mutation rate in L1-derived tissues compared to other somatic cell types^13,87^. Although to a lesser extent than the epidermis, this interesting contrast, where the cell-type restricted TF motifs are conserved and the cell-type-specific chromatin accessibility of cognate ACRs are not, extends to other cell types. This supports a larger pattern of novel ACR evolution that co-opts established cell-type-specific TF networks.

**Fig. 7.**
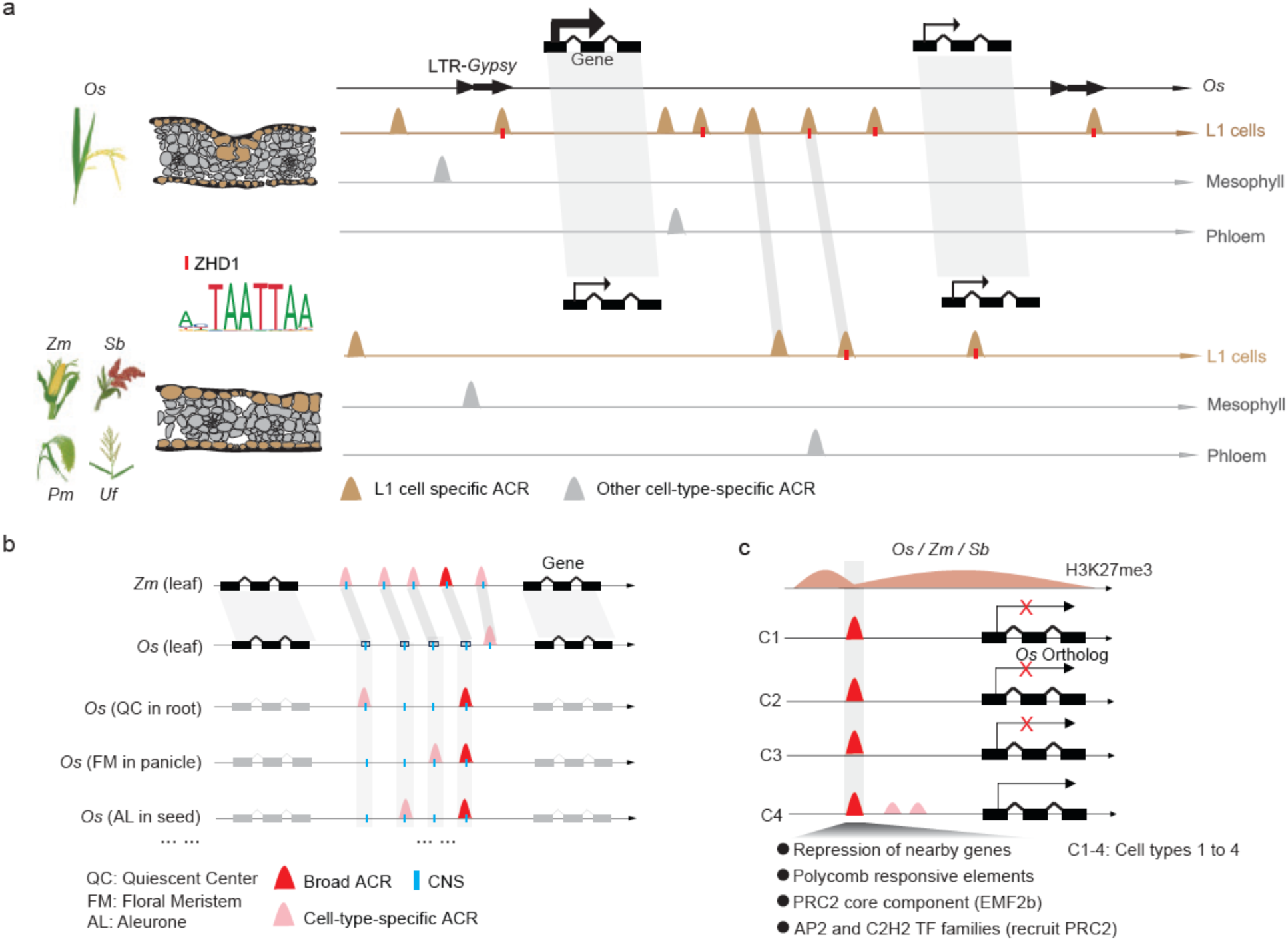
Evolution of cell-type-specific ACRs and CREs. **a,** The analysis of leaf cell types across these species revealed an enrichment of cell-type-specific ACRs in species-specific regions. Notably, these species-specific ACRs were enriched within L1-derived cells compared to all others examined. Additionally, it was observed that epidermal cell-specific ACRs significantly overlapped with LTR-*Gypsy* TEs, which were enriched for the ZHD1 motif known to regulate leaf curling. The L1 ACRs within the ZHD1 motif are likely associated with species-specific elevated expression of a small number of genes involved in L1-derived cell development. **b,** We found an enrichment of CNS in cell-type-specific ACRs. Although some CNS ACRs retained the same cell-type specificity between *O. sativa* and *Z. mays,* these CNS ACRs often switched tissue or cell-type accessibility between grass species. **c,** Despite being within facultative heterochromatin, H3K27me3-broad ACRs are accessible in many cell types, providing a physical entry point for PRC2 to bind. Several lines of evidence support that the H3K27me3-broad ACRs contain silencer CREs. Specifically, these ACRs are linked to transcriptionally silent genes, are enriched for PRE motifs, enriched for TF family motifs (AP2 and C2H2) reported to recruit PRC2, and enriched for PRC2 subunit (EMF2b) ChIP-seq peaks.

Highlighting the rapid rate of regulatory evolution, ACRs, and the CREs within them, underpin phenotypic variation within plant and mammalian species^10,20,88,89^. Despite the link between CREs and phenotypic variation, how these transcriptional regulatory circuits have changed during species divergence is challenging to address. This is partly due to rapid CRE changes occluding pairwise comparison; even closely related plant species share but a fraction of their ACR/CRE complements^6,90^. We used a comparative single-cell epigenomics approach to characterize the evolution of cell-type-specific ACRs and CREs in grasses. We demonstrate that grass cell-type-specific ACRs have changed tremendously over 50 million years^58^, with relatively few (0.7% to 1.8%) cell-type-specific ACRs remaining conserved across the examined plant species (Extended Data Fig. 5b and f). This contrasts with mammals, where previous work on liver specific enhancers found that ∼5% (2,151/43,020) remain conserved over ∼100 million years^91^. The difference in ACR conservation highlights the speed at which plant CRE evolution takes place compared to mammals, which has been supported previously^59,90,92-95^. The repeated whole genome duplications in plant lineages^96,97^, and the functional redundancy they provide, may be the fuel for rapid CRE divergence driving plants’ adaptation to diverse environments^98^.

Integration of the *O. sativa* atlas with CNS revealed ∼65% were accessible in at least one cell type (Fig. 5a). We expect that most CNS not captured by an ACR in our study are likely accessible in an unsampled cell type, environmental, or developmental condition. This stresses the need for expanded accessible chromatin atlases using more tissues, segments of development, and environmental conditions. Most ACRs containing CNS had variable cell-type specificity between species (Fig. 7b), highlighting that deeply conserved grass ACRs readily evolve new spatiotemporal usage. Although less frequently than grass ACRs, around one third of conserved mouse DNAseI–hypersensitive sites (DHSs) are altered in human tissue contexts after ∼90 million years of evolution^69^. Thus, although eukaryotic CNS exhibit sequence conservation, their functional context is often altered^71^, with novel spatiotemporal CNS usage appearing prevalent in grass lineages. However, it remains possible that the main CNS function conserved between *O. sativa* and *Z. mays* occurs in non-leaf tissues. Nonetheless, the switching of cell-type accessibility highlights the importance of merging chromatin accessibility data with CNS datasets, as the assumption of conserved CNS sequence equaling conserved CRE function is often invalid.

Much focus has been placed on enhancer CREs within ACRs, yet silencers are equally important, as they repress gene expression until the proper developmental or environmental cues. Our prior research uncovered that some ACRs flanking H3K27me3 are linked to the suppression of nearby genes^6,9,20^; however, the question remains whether these ACRs function as silencers. Our *O. sativa* cell-type ACR atlas allowed the identification of ACRs within H3K27me3 regions that were accessible in most cell types. Several lines of evidence support these H3K27me3-broad ACRs as silencers of linked, transcriptionally repressed genes. Specifically, these putative silencers were enriched for CNS, PREs and related TF motifs, and PRC2 *in vivo* occupancy^83^ (Fig. 7c). Additional comparison of two genotypes in *O. sativa* reveals the PRE motif mutations within H3K27me3-broad ACRs may preclude PRC2 targeting (Extended Data Fig. 10f), but future genome editing of these putative silencers will reveal more about their role in PRC2 recruitment and gene silencing. Similar H3K27me3-broad ACR putative silencers were also present in the epigenomic landscape of *O. sativa*, *Z. mays*, and *S. bicolor*, suggesting this is a conserved feature of grass genomes. H3K27me3 silencing is deeply conserved in eukaryotes^99^, and a recent study found that many H3K27me3-marked regions might function as silencer-like regulatory elements in *O. sativa*^76^. We hypothesize that other single-cell comparative genomic investigations will find this pattern of broadly accessible silencers in other angiosperm species with H3K27me3.

Our rice atlas of cell-type-specific ACRs and this cross-species analysis provides a useful resource to enhance our understanding of regulatory evolution more broadly. Highlighting the rapid rate of regulatory evolution, we believe combining this data with that from more closely related grasses in the future will reveal more nuanced evolutionary dynamics of CRE evolution under different levels of evolutionary time. This resource, and these observations, will fuel research into identifying key CREs controlling specific genes by demarcating high-confident targets for genome editing.

## Methods

### Preparation of plant materials

Early seedlings, specifically seedling tissues above ground, were collected 7 and 14 days after sowing. Flag leaf tissue was harvested at the V4 stage, characterized by collar formation on leaf 4 of the main stem. Axillary buds were obtained from rice plants grown in the greenhouse at approximately the V8 stage. Rice seminal and crown root tips (bottom 2 cm) were gathered at the same stage as seedling tissues, 14 days after sowing. Panicle tissue was acquired from rice plants grown in the greenhouse. Inflorescence primordia (2-5 mm) were extracted from shoots harvested at the R1 growth stage, where panicle branches had formed. Early seeds were harvested at approximately six days after pollination (DAP), and late seeds at approximately ten DAP. All tissues were collected between 8 and 9 am, and all fresh materials were promptly utilized for library construction starting at 10 am.

### Single-cell ATAC-seq library preparation

Nuclei isolation and purification were performed as described previously^100^. In brief, the tissue was finely chopped on ice for approximately 2 minutes using 600 μL of pre-chilled Nuclei Isolation Buffer (NIB, 10 mM MES-KOH at pH 5.4, 10 mM NaCl, 250 mM sucrose, 0.1 mM spermine, 0.5 mM spermidine, 1 mM DTT, 1% BSA, and 0.5% TritonX-100). After chopping, the entire mixture was passed through a 40-μm cell strainer and then subjected to centrifugation at 500 rcf for 5 minutes at 4°C. The supernatant was carefully decanted, and the pellet was reconstituted in 500 μL of NIB wash buffer, which consisted of 10 mM MES-KOH at pH 5.4, 10 mM NaCl, 250 mM sucrose, 0.1 mM spermine, 0.5 mM spermidine, 1 mM DTT, and 1% BSA. The sample was filtered again, this time through a 10-μm cell strainer, and then gently layered onto the surface of 1 mL of a 35% Percoll buffer, prepared by mixing 35% Percoll with 65% NIB wash buffer, in a 1.5-mL centrifuge tube. The nuclei were subjected to centrifugation at 500 rcf for 10 minutes at 4°C. Following centrifugation, the supernatant was carefully removed, and the pellets were resuspended in 10 μL of diluted nuclei buffer (DNB, 10X Genomics Cat# 2000207). About 5 μL of nuclei were diluted 10 times and stained with DAPI (Sigma Cat. D9542) and then the nuclei quality and density were evaluated with a hemocytometer under an epifluorescence microscope. The original nuclei were diluted with a DNB buffer to a final concentration of 3,200 nuclei per uL. Finally, 5 uL of nuclei (16,000 nuclei in total) were used as input for scATAC-seq library preparation. scATAC-seq libraries were prepared using the Chromium scATAC v1.1 (Next GEM) kit from 10X Genomics (Cat# 1000175), following the manufacturer’s instructions. (10xGenomics, CG000209_Chromium_NextGEM_SingleCell_ATAC_ReagentKits_v1.1_UserGuide_RevE). Libraries were sequenced with Illumina NovaSeq 6000 in dual-index mode with eight and 16 cycles for i7 and i5 index, respectively.

### Single-nuclei RNA-seq library preparation and data analysis

The protocol for nuclei isolation and purification was adapted from the previously described scATAC-seq method. In summary, to minimize RNA degradation and leakage, the tissue was finely chopped on ice for approximately 1 minute using 600 μL of pre-chilled Nuclei Isolation Buffer containing 0.4U/μL RNase inhibitor (Roche, Protector RNase Inhibitor, Cat. RNAINH-RO) and a comparatively low detergent concentration of 0.1% NP-40. Following chopping, the entire mixture was passed through a 40-μm cell strainer and then subjected to centrifugation at 500 rcf for 5 minutes at 4°C. The supernatant was carefully decanted, and the pellet was reconstituted in 500 μL of NIB wash buffer, comprising 10 mM MES-KOH at pH 5.4, 10 mM NaCl, 250 mM sucrose, 0.5% BSA, and 0.2U/μL RNase inhibitor. The sample was filtered again, this time through a 10-μm cell strainer, and gently layered onto the surface of 1 mL of a 35% Percoll buffer. The Percoll buffer was prepared by mixing 35% Percoll with 65% NIB wash buffer in a 1.5-mL centrifuge tube. The nuclei were then subjected to centrifugation at 500 rcf for 10 minutes at 4°C. After centrifugation, the supernatant was carefully removed, and the pellets were resuspended in 50 μL of NIB wash buffer. Approximately 5 μL of nuclei were diluted tenfold and stained with DAPI (Sigma Cat. D9542). Subsequently, the nuclei’s quality and density were evaluated with a hemocytometer under a microscope. The original nuclei were further diluted with a DNB buffer to achieve a final concentration of 1,000 nuclei per μL. Ultimately, a total of 16,000 nuclei were used as input for snRNA-seq library preparation. For snRNA-seq library preparation, we employed the Chromium Next GEM Single Cell 3’GEM Kit v3.1 from 10X Genomics (Cat# PN-1000123), following the manufacturer’s instructions (10xGenomics, CG000315_ChromiumNextGEMSingleCell3-_GeneExpression_v3.1_DualIndex_RevB). The libraries were subsequently sequenced using the Illumina NovaSeq 6000 in dual-index mode with 10 cycles for the i7 and i5 indices, respectively.

The raw BCL files obtained after sequencing were demultiplexed and converted into FASTQ format using the default settings of the 10X Genomics tool cellranger mkfastq^101^ (v7.0.0). The raw reads were processed with cellranger count^101^ (v7.0.0) using the Japonica rice reference genome^102^ (v7.0). Genes were kept if they were expressed in more than three cells and each cell having a gene expression level of at least 1,000 but no more than 10,000 expressed genes. Cells with over 5% mitochondria or chloroplast counts were filtered out. The expression matrix was normalized to mitigate batch effects based on global-scaling normalization and multicanonical correlation analysis in Seurat^103^ (v4.0). The Scrublet tool^104^ was employed to predict doublet cells in this dataset. SCTransform in Seurat was used to normalize the data and identify variable genes. The nearest neighbors were computed using FindNeighbors using 30 PCA dimensions. The clusters were identified using FindClusters with a resolution of 1. The cell types were annotated based on the marker gene list (Supplementary Table 2). To identify genes exhibiting higher expression in a particular cell type than in the others, we utilized the ‘cpm’ function from edgeR^105^ (v3.38.1), for normalizing the expression matrix. Genes within a specific cell type that displayed more than 1.5-fold change in log2 Counts Per Million (CPM) values compared to the average log2 CPM across all cell types were determined as specifically expressed genes in that particular cell type.

Transcriptome similarity between cell types of *O. sativa* and *Z. mays* was assessed using the MetaNeighbor package^106^. To statistically compare similarities across different cell types, we randomly divided the cells of each type into five groups. Each group was then used as input for the MetaNeighbor analysis. The area under the receiver operating characteristic curve (auROC) score obtained from MetaNeighbor was used as the similarity score in our analysis.

### Slide-seq library preparation and data analysis

Root tissues from rice seedlings 14 days after sowing were used for the Slide-seq V2 spatial transcriptomics. The tissues were embedded in the Optimal Cutting Temperature (OCT) compound, snap-frozen in a cold 2-methylbutane bath, and cryosectioned into 10 um thick slices. The spatial transcriptome library was constructed following a published method^107,108^. In brief, the tissue slices were placed on the Slide-seq V2 puck and underwent the RNA hybridization and reverse transcription process. After tissue clearing and spatial bead collections, the cDNA was synthesized and amplified for a total of 14 cycles. The library was constructed using Nextera XT Library Prep Kit (Illumina, USA) following the manufacturer’s instructions.

The reads alignment and quantification were conducted following the Slide-seq pipeline (https://github.com/MacoskoLab/slideseq-tools). The data processing is similar to the procedures applied in the analysis of snRNA-seq but setting a resolution of 0.7 for the FindClusters function in Seurat^103^ (v4.0). The cell types were annotated based on the histology of cross-sectioned roots. The marker genes of each cluster were identified using the Wilcoxon signed rank test in FindAllMarkers.

### RNA *in situ* Hybridization

The rice samples were put into the vacuum tissue processor (HistoCore PEARL, Leica) to fix, dehydrate, clear, and embed, and were subsequently embedded in paraffin (Paraplast Plus, Leica). The samples were sliced into 8 μm sections with a microtome (Leica RM2265). The cDNAs of the genes were amplified with their specific primer pairs in situ_F/in situ_R and subcloned into the pGEM-T vector (Supplementary Table 27). The pGEM-gene vectors were used as the template to generate sense and antisense RNA probes. Digoxigenin-labeled RNA probes were prepared using a DIG Northern Starter Kit (Roche) according to the manufacturer’s instructions. Slides were observed under bright fields through a microscope (ZEISS) and photographed with an Axiocam 512 color charge-coupled device (CCD) camera.

### Raw reads processing of scATAC-seq

Data processing was executed independently for each tissue and/or replicate. Initially, raw BCL files were demultiplexed and converted into FASTQ format, utilizing the default settings of the 10X Genomics tool cellranger-atac make-fastq^109^ (v1.2.0). Employing the Japonica rice reference genome^102^ (v7.0), the raw reads underwent processing using cellranger-atac count^109^ (v1.2.0). These steps encompassed adaptor/quality trimming, mapping, and barcode attachment/correction. Subsequently to the initial processing, reads that were uniquely mapped with mapping quality > 10 and correctly paired were subjected to further refinement through SAMtools view (v1.7; -f 3 -q 10; Li et al. 2009). To mitigate the impact of polymerase chain reaction (PCR) duplicates, the Picard tool MarkDuplicates^110^ (v2.16.0) was applied on a per-nucleus basis. To elevate data quality, a blacklist of regions was devised to exclude potentially spurious reads. The methodology involved the exclusion of regions displaying bias in Tn5 integration from Tn5-treated genomic DNA. Specifically, regions characterized by 1-kilobase windows with coverage exceeding four times the genome-wide median were eliminated. We further leveraged ChIP-seq input data^6^, to filter out collapsed sequences in the reference using the same criteria. This blacklist also incorporated sequences of low-complexity and homopolymeric sequences through RepeatMasker^111^ (v4.1.2). Moreover, nuclear sequences exhibiting homology surpassing 80% to mitochondrial and chloroplast genomes^112^ (BLAST+; v2.11.0) were also included within the blacklist. Furthermore, BAM alignments were converted into BED format, wherein the coordinates of reads mapping to positive and negative strands were subjected to a shift by +4 and -5, respectively. The unique Tn5 integration sites per barcode were finally retained for subsequent analyses.

### Identifying high-quality nuclei

To ensure the acquisition of high-quality nuclei, we harnessed the capabilities of the Socrates package for streamlined processing^20^. To gauge the fraction of reads within peaks, we employed MACS2^113^ (v2.2.7.1) with specific parameters (genomesize = 3e8, shift= -75, extsize = 150, fdr = 0.05) on the bulk Tn5 integration sites. Subsequently, we quantified the number of integration sites per barcode using the callACRs function. Next, we estimated the proximity of Tn5 integration sites to genes, focusing on a 2 kb window surrounding the TSS. This estimation was achieved through the buildMetaData function, which culminated in the creation of a meta file. For further refinement of cell selection, we harnessed the findCells function, implementing several criteria: 1) A minimum read depth of 1,000 Tn5 integration sites was required. 2) The total number of cells was capped at 16,000. 3) The proportion of reads mapping to TSS sites was above 0.2, accompanied by a z-score threshold of 3. 4) Barcode FRiP scores were required to surpass 0.1, alongside a z-score threshold of 2. 5) We filtered out barcodes exhibiting a proportion of reads mapping to mitochondrial and chloroplast genomes that exceeded two standard deviations from the library mean. 6) We finally used the detectDoublets function to estimate doublet likelihood by creating synthetic doublets and conducting enrichment analysis. These multiple steps ensured the meticulous identification and selection of individual cells, facilitating a robust foundation for subsequent analyses.

### Nuclei clustering

For the nuclei clustering, we leveraged all functions from the Socrates package^20^. We binned the entire genome into consecutive windows, each spanning 500 bp. We then tabulated the count of windows featuring Tn5 insertions per cell. Barcodes falling below one standard deviation from the mean feature counts (with a z-score less than 1) were excluded. Moreover, barcodes with fewer than 1,000 features were eliminated. We pruned windows that exhibited accessibility in less than 0.5% or more than 99.5% of all nuclei. To standardize the cleaned matrix, we applied a term frequency-inverse document frequency normalization function. The dimensionality of the normalized matrix underwent reduction through the utilization of non-negative matrix factorization, facilitated by the R package RcppML^114^ (v0.3.7). We retained 50 column vectors from an uncentered matrix. Subsequently, we selected the top 30,000 windows that displayed the highest residual variance across all cells. This selection was based on fitting a model where the proportion of cells with accessibility served as an independent variable and variance as the dependent variable. To further reduce the dimensionality of the nuclei embedding, we employed the UMAP technique using umap-learn (k = 50, min_dist = 0.1, metrix = ‘euclidean’) in R^115^ (v0.2.8.0). Furthermore, we clustered nuclei using the callClusters function within the Socrates framework^20^. Louvain clustering was applied, with a setting of k=50 nearest neighbors at a resolution of 0.7. This process underwent 100 iterations with 100 random starts. Clusters with an aggregated read depth of less than 1 million and 50 cells were subsequently eliminated. To filter outlying cells in the UMAP embedding, we estimated the mean distance for each nucleus using its 50 nearest neighbors. Nuclei that exceeded 3 standard deviations from the mean distance were deemed outliers and removed from consideration.

### Estimation of gene accessibility scores

To estimate the gene accessibility, we employed a strategy wherein the Tn5 insertion was counted across both the gene body region as well as a 500 bp extension upstream. Subsequently, we employed the SCTransform algorithm from the Seurat package^103^ (v4.0) to normalize the count matrix that was then transformed into a normalized accessibility score, with all positive values scaled to 1.

### Cell type validation

Upon completing the initial annotation process, which was based on a curated list of marker genes, we further expanded our marker repertoire by incorporating markers collected from published bulk RNA-seq data encompassing a diverse array of cell types, which were acquired *via* laser capture dissection, as well as a published scRNA-seq data encompassing panicle related cell types^116^. In brief, we collected the markers from several studies, including three distinct cell types within rice leaves^117^ and ten cell types across rice seed organs^118,119^. From these sources, we selected the top 100 variably expressed markers for each cell type and employed them to compute cell identity enrichment scores. We undertook a comprehensive assessment of markers linked to the different cell types. Subsequently, we randomly drew 100 markers from this pool and repeated this procedure 1,000 times to construct a null distribution based on marker chromatin accessibility scores. For each target cell type marker, we compared their accessibility scores to this null distribution. This facilitated the derivation of an enrichment score per cell, delineating the marker’s significance for each representative cell type. We next employed a MAGIC algorithm^120^ to refine these enrichment scores. These scores were then mapped onto a UMAP plot, enhancing the cell identity annotation.

Furthermore, we undertook validation of our single-cell chromatin accessibility atlas through integration with published scRNA-seq from rice root tissue. This validation was achieved through two distinct approaches (Supplementary Fig. 8). In the first approach, we leveraged the marked enrichment technique, adapting the above mentioned methodology with the incorporation of the top 20 markers derived from marker identification using the ‘FindMarkers’ function in Seurat^103^ (v4.0). Following the acquisition of a smoothed score for each cell type, individual cells were annotated to specific cell types based on the largest enrichment score within that cell type. A threshold was further set, requiring the maximum score to exceed 0.5 for confident labeling; otherwise, the cell was labeled as ‘Unknown’. The second approach entailed employing a k-nearest neighbor (knn) strategy. This strategy commenced with the normalization of scRNA-seq datasets, mirroring the process applied to scATAC-seq datasets. The top 3,000 most variable genes within the scATAC-seq dataset were then identified using the Seurat function ‘FindVariableFeatures’, subsequently filtering to include only genes common to both datasets. By treating the scRNA-seq cells as a reference, a dimension reduction process was conducted to generate a loading matrix, which was then utilized to project the scATAC-seq cells onto the scRNA-seq cell embedding. The integration of these two datasets was achieved through the Harmony algorithm^121^ (v0.1.0). Within the dual embeddings, the 20 nearest neighbors of each scATAC-seq cell in the scRNA-seq dataset were computed. The most frequent label among these RNA neighbors (> 10 cells) was subsequently assigned as the label for each scATAC-seq cell or designated as NA if no label meeting this threshold was identified.

### Identification of *de novo* marker genes

For each cell type, we used edgeR^105^, to identify cell-type-specific genes that are differentially accessible. To determine if genes are accessible in a specific cell type, we compared the genes in the target cell type to those in all other cell types, which served as the reference. We identified genes with an FDR < 0.05 and log2 (Fold change) > 0 as candidate genes that are specifically accessible in a particular cell type.

### Cell cycle prediction

The prediction of cell cycle stages per nucleus was executed similarly to annotating cell identities based on the aforementioned enriched scores. In brief, we collected a set of 55 cell-cycle marker genes from a previous study^122^. For every cell-cycle stage, the cumulative gene accessibility score for each nucleus was computed. These resultant scores were subsequently normalized using the mean and standard deviation derived from 1,000 permutations of the 55 random cell-cycle stage genes, with exclusion of the focal stage. Z-scores corresponding to each cell-cycle stage were transformed into probabilities using the ‘pnorm’ function in R. Furthermore, the cell-cycle stage displaying the highest probability was designated as the most probable cell stage.

### ACR identification

Upon segregating the comprehensive single-base resolution Tn5 insertion sites BED dataset into distinct subsets aligned with annotated cell types, we executed the MACS2 tool^113^ (v2.2.7.1) for precise peak identification per cell type. Notably we employed non-default parameters, specifically: --extsize 150, --shift -75, -nomodel -keep-dup all. To mitigate potential false positives, a permutation strategy was applied, generating an equal number of peaks based on regions that were mappable and non-exonic. This approach encompassed the assessment of Tn5 insertion sites and density within both the original and permuted peak groups. By scrutinizing the permutation outcomes, we devised empirically derived false discovery rate (FDR) thresholds specific to each cell type. This entailed determining the minimum Tn5 density score within the permutation cohort where the FDR remained < 0.05. To further eliminate peaks that exhibited significant overlap with nucleosomes, we applied the NucleoATAC tool^123^ (v0.2.1) to identify potential nucleosome placements. Peaks that featured over 50% alignment with predicted nucleosomes were systematically removed. The average fragment size of reads overlapping with the peaks were calculated and the peaks with the average fragment size > 150 bp were filtered out. Ultimately, the pool of peaks for each cell type was amalgamated and fine-tuned, yielding 500 bp windows that were centered on the summit of ACR coverage.

### Identification of cell-type-specific ACRs

To identify cell-type-specific ACRs across the *O. sativa* atlas, we used a modified entropy metric in our prior study^24^. Briefly, this method calculates the accessibility of ACRs, normalizes these values using CPM, and then calculates both the entropy and specificity scores for each ACR in each cell type. We combined this method with a bootstrapping approach^124^. For each cell type and species, a sample of 250 cells was taken 5,000 times with replacement, and the specificity metric was calculated as described above. This specificity metric was then compared against a series of 5,000 null distributions, each consisting of a random shuffle of 250 cells from mixed cell populations. This allowed us to set thresholds for what we would expect from non-cell-type-specific data. A nonparametric test was then used to compare the median real bootstrap specificity score to the null distributions. ACRs were labeled as cell-type-specific if they had a *p* value less than 0.001. Cell-type-specific ACRs were those with a significant *p* value in a given cell type and found to be specific in one or two cell types for leaf ACRs across five examined species, and in one, two or three cell types for ACRs in the *O. sativa* atlas. This method was used for all species in this study. Due to the number of ACRs and cell types in the *O. sativa* atlas, each tissue was analyzed independently, and the results were merged downstream.

### Interactions between distal cell-type-specific ACRs and cell-type-specific genes

Raw Hi-C data from *O. sativa* were collected including five replicates^25^. Low-quality reads were filtered out using Trimmomatic^125^. The clean reads from each replicate were mapped to the Japonica rice reference genome^102^. These mapped reads were then used to obtain normalized contact maps through a two-step approach in the HiC-Pro software^126^ in parallel mode. The analysis was run using the default configuration file, with modifications to specify a minimal mapping quality of 10 and enzyme recognition sites of MboI. The valid pairs files obtained from all replicates of the same species were combined using the ‘merge_persample’ step in HiC-Pro for further analysis. Fit-HiC^127^ was used to identify intra-chromatin loops. The input contact maps for Fit-HiC were generated from valid pairs files using the ‘validPairs2FitHiC-fixedSize’ script at a 5 kb resolution. Fragments and bias files were generated using ‘creatFitHiCFragments-fixedsize’ and ‘HiCKRy’ scripts separately. Significant intra-chromosomal interactions were identified by running FitHiC at a 5 kb resolution. These significant interactions were further merged and filtered using ‘merge-filter.sh’ script at a 5 kb resolution and with a FDR of 0.05. Each bin of the significant intra-chromatin loops was intersected with distal cell-type-specific ACRs using the ‘intersect’ function in bedtools^128^. The bins associated with those intersecting the distal cell-type-specific ACRs were further examined to determine if any intersected with promoter regions of cell-type-specific genes, defined as regions 2 kb upstream of the genes.

### Aligning pseudotime trajectories between rice and maize

Motif deviation scores were computed utilizing ChromVAR^129^ (v1.18.0) for both rice and maize cells originated from the trajectories of root developing xylem and cortex development. To refine these scores, a diffusion approach based on the MAGIC algorithm^130^ was used. We further used cellAlign tool^131^ (v0.1.0), a technique that standardized the imputed deviation scores across a predefined set of points (n = 200) distributed evenly along the trajectories of rice and maize. This normalization strategy aimed to mitigate technical biases inherent in the data. For each motif pair shared between rice and maize, a comprehensive global alignment procedure ensured to align the imputed deviation scores across pseudotime for both *O. sativa* and *Z. mays*. Subsequently, we calculated the normalized distance between the two species using the cellAlign tool. Motif clustering based on these distance scores, using k-means, yielded two distinct groups: Group 1, characterized by relatively higher distances, and Group 2. A linear regression framework was subsequently introduced, using distance scores and pseudotime as predictive variables for motifs within these two groups. In instances where motif pairs within either group exhibited positive or negative coefficients, we classified them as ‘shiftEarlyrice’ or ‘shiftEarlymaize’. Notably, motif pairs in Group 1 were designated as ‘conserved’ if their coefficients bore identical positive or negative attributes. To enhance the robustness of our findings, *p* values acquired from the linear regression analysis underwent adjustment using the Benjamini-Hochberg procedure, effectively addressing multiple comparisons. If the corrected *p* value exceeded 0.05, the motif pairs were categorized as ‘Unknown’.

### Correlation between chromatin accessibility of TF genes and motif deviatio**n**

We sourced rice and *A. thaliana* TFs from PlantTFDB^132^ (v4.0) database. To identify rice orthologs of *A. thaliana* TFs, we employed BLAST^112^ (BLAST+; v2.11.0) by utilizing protein fasta alignments with an e-value threshold of 1e-5 used for significance. Alignments were restricted to fasta sequences categorized as TFs from either species. To further refine the putative orthologs, we applied filters based on functional similarity to *A. thaliana* TFs. Alignments with less than 15% identity were excluded, along with rice TFs associated with distinct families. From the remaining candidates, we selected the orthologs demonstrating the highest Pearson correlation coefficient concerning the motif deviation scores. Motif deviation scores of specific TF motifs within nuclei were computed *via* chromVAR^129^ (v1.18.0).

### Linear-model based motif enrichment analysis

We employed the FIMO tool from the MEME suite^133^ (v5.1.1) with a significance threshold of *p* value < 10^-5^ to predict motif locations. The motif frequency matrix used was sourced from the JASPAR plants motif database^134^ (v9). Subsequently, we constructed a binarized peak-by-motif matrix and a motif-by-cell count matrix. This involved multiplying the peak-by-cell matrix with the peak-by-motif matrix. To address potential overrepresentation and computational efficiency, down-sampling was implemented. Specifically, we standardized the cell count by randomly selecting 412 cells per cell type per species. This count represents the lowest observed cell count for a given cell type across all species. For each cell type annotation, total motif counts were predicted through negative binomial regression. This involved two input variables: an indicator column for the annotation, serving as the primary variable of interest, and a covariate representing the logarithm of the total number of nonzero entries in the input peak matrix for each cell. The regression provided coefficients for the annotation indicator column and an intercept. These coefficients facilitated the estimation of fold changes in motif counts for the annotation of interest in relation to cells from all other annotations. This iterative process was conducted for all motifs across all cell types. The obtained *p* values were adjusted using the Benjamini-Hochberg procedure to account for multiple comparisons. Finally, enriched motifs were identified by applying a dual filter criterion: corrected *p* values < 0.01, fold-change of the top enriched TF motif in cell type-specific peaks for all cell types should be over 1, and beta (motif enrichment score) > 0.05 or beta > 0.

### Binomial test-based motif enrichment analysis

To assess the enrichment of motifs in a target set of ACRs, we performed analysis for each specific motif. We randomly selected an equivalent number of ACRs as found in the target set, repeating this process 100 times. Notably, the randomly selected ACR set did not overlap with the actual target set of ACRs. Following this, we computed the average ratio of ACRs capturing the motif within the null distribution.

Subsequently, we executed an exact Binomial test^135^, wherein we set this ratio as the hypothesized probability of success. The number of ACRs overlapping the motif in the target set was considered the number of successes, while the total number of ACRs in the target set represented the number of trials. The alternative hypothesis was specified as ‘two.sided’. This meticulous approach allowed us to robustly evaluate and identify significant motif enrichments within the target set of ACRs.

### Construction of control sets for enrichment tests

To check if non-CDS QTNs could be significantly captured by ACRs, we generated control sets by simulating sequences with the same length as non-CDS QTNs 100 times, yielding a mean proportion for the control sets. The binomial test *p* value was calculated by comparing the mean ratio to the observed overlapping ratio of non-CDS QTNs captured by ACRs.

To check if ACRs could significantly capture CNS, we generated control sets by simulating sequences with the same length as ACRs 100 times, yielding a mean proportion for the control sets. The binomial test *p* value was calculated by comparing the mean ratio to the observed overlapping ratio of ACRs capturing the CNS.

To perform comparative analysis of expression levels and chromatin accessibility of genes surrounding broad ACRs under and outside of H3K27me3 peaks, we sampled the same number of ACRs per cell type regarding the broad ACRs not under H3K27me3 peaks. This step is to make sure that their nearby gene chromatin accessibility exhibited similar values compared to the broad ACRs under the H3K27me3 peaks.

To check if the H3K27me3-broad-ACRs could significantly capture the known PREs and capture the EMF2b ChIP-seq peaks, we generated control sets by randomly selecting not-H3K27me3-broad-ACR instances 100 times, yielding a mean number value for the control sets. The Binomial test *p* value was calculated by comparing the mean ratio to the observed number of H3K27me3-broad ACRs overlapped with the PREs.

To test if H3K27me3-broad ACRs in *O. sativa*, *Z. mays*, and *S. bicolor* significantly capture six known motifs, we generated control sets by simulating sequences with the same length as ACRs 100 times, yielding a mean proportion for the control sets. The binomial test *p* value was calculated by comparing the mean ratio to the observed overlapping ratio of H3K27me3-broad ACRs capturing the motifs. The same process was conducted to examine whether the EMF2b peaks significantly capture the six motifs.

### Identification of H3K27me3-broad ACRs

We first implemented a series of cutoffs to determine whether the peak is accessible for a specific cell type. For each cell type, we first normalized the read coverage depth obtained from the MACS2 tool divided by total count of reads, and ensured that the maximum of normalized coverage within the peak exceeded a predefined threshold set at 2. Additionally, we calculated Tn5 integration sites per peak, filtering out peaks with fewer than 20 integration sites. Subsequently, we constructed a peak by cell type matrix with Tn5 integration site counts. This matrix underwent normalization using the ‘cpm’ function wrapped in edgeR^105^ (v3.38.1) and ‘normalize.quantiles’ function wrapped within preprocessCore^136^ (v1.57.1) in the R programming environment. To further refine our selection, a threshold of 2 was set for the counts per million value per peak per cell type. Peaks that satisfied these distinct cutoff criteria were deemed accessible in the designated cell types. For analyzing cell types with fewer than 10 samples (n < 10), we established criteria where H3K27me3-broad ACRs must be accessible in at least n-1 cell types, while cell-type-specific ACRs should be accessible in fewer than 3 of the examined cell types. For analyses involving more than 10 cell types, we adjusted the criteria: H3K27me3-broad ACRs must be accessible in at least n−2 cell types, and cell-type-specific ACRs should be accessible in fewer than 4 of the examined cell types.

### *De novo* motif analysis

To identify position weight matrix of six known motifs within 170 *A. thaliana* PREs^74^, we employed the streme function with default settings from the MEME suite^133^ (v5.1.1). The control sequences were built up to match each PRE sequence by excluding exons, PREs, and unmappable regions, and they possess a similar GC content (< 5% average difference) and same sequence length compared to the positive set.

### Identification of syntenic regions

Identification of syntenic gene blocks was done using the GENESPACE^137^ (v1.4). In brief, to establish orthologous relationships between ACR sequences, ACRs in the *O. sativa* genome were extended to incorporate the two closest gene models for a ‘query block’ since GENESPACE only draws relationships between protein coding sequences. Then the GENESPACE function ‘query_hits’ was used with the argument ‘synOnly = TRUE’ to retrieve syntenic blocks. The resulting syntenic hits were further filtered to allow only a one-to-one relationship between *O.sativa* and the corresponding species. The corresponding syntenic blocks were then named and numbered, and both the genes and genomic coordinates were recorded.

To further identify corresponding ACRs within these blocks we set up a BLASTN pipeline^138^ (v2.13.0). For each comparison of species, using *O. sativa* as the reference the underlying nucleotide sequences of the syntenic regions were extracted using Seqkit, and used as the blast reference database^139^ (v2.5.1). The sequences underlying the ACRs within the same syntenic region in a different species were then used as the query. The blast was done using the following parameters to allow for alignment of shorter sequences ‘-task blastn-short -evalue 1e-3 - max_target_seqs 4-word_size 7 -gapopen 5 -gapextend 2 -penalty -1 -reward 1 -outfmt 6’. This procedure was run for each syntenic region separately for all species comparisons. The resulting BLASTN files were combined, and then filtered using a custom script. Alignments were only considered valid if the e-value passed a stringent threshold of 1e-3, and the alignment was greater than 20 nucleotides with the majority of the shared ACRs (92% to 94%) containing the alignment regions including TF motif binding sites (Supplementary Fig. 19). The resulting filtered BLAST files, and the BED files generated from these BLAST files allowed us to draw our relationships between ACRs in the corresponding syntenic space. For all analyses, ACRs were considered to have conserved cell-type-specificity if these ACRs would be aligned by BLAST and had the same cell-type as assigned by the above method.

### Estimation of conservation scores

Conservation scores were predicted using PhyloP^140^ (v1.0), where values are scaled between 0 to 1, with one being highly conserved and 0 being non-conserved. Phylogenies to train PhyloP were generated using PhyloFit^141^ (v1.0), and neutral and conserved sequences were identified using the whole genome aligner progressive cactus.

### ChIP-seq analysis

The clean reads of EMF2b were downloaded from a previous study^83^. The reads were mapped to the rice reference genome^102^ (v7.0) using bowtie2^142^ (v2.5.2) with the following parameters: ‘-- very-sensitive --end-to-end’. Reads with MAPQ > 5 were used for the subsequent analysis. Aligned reads were sorted and duplicated reads were removed using SAMtools^143^ (v1.7). Peak calling was performed using epic2^144^ with the following parameters: ‘-fdr 0.01 --bin-size 150 -- gaps-allowed 1’. The peak ‘BED’ and ‘BIGWIG’ files of H3K27me3 ChIP-seq data for leaf, root, and panicle rice organs were downloaded from RiceENCODE^145^ (http://glab.hzau.edu.cn/RiceENCODE/).

### Genetic variants calling in ZS97 genotype in *O. sativa*

We obtained raw sequencing data for the ZS97 genotype of *O. sativa* from a published study^85^. After quality filtering the raw reads using fastp^146^ (v0.23.4), we aligned them to the Japonica *O. sativa* reference genome (Ouyang et al., 2007) using the BWA-MEM algorithm^147^ (v0.7.8). We then used Picard tool MarkDuplicates^110^ (v2.16.0) to remove PCR duplicates. The final genetic variants file was generated using the HaplotypeCaller function in GATK^148^ (GATK 4.2.3.0).

### GO enrichment test

The GO enrichment tests were performed based on the AgriGO^149^ (v2) by setting the Chi-square statistical test and multi-test adjustment method is Hochberg (FDR).

## Additional resources

Cell-type resolved data can be viewed through our public Plant Epigenome JBrowse Genome Browser^150^ (http://epigenome.genetics.uga.edu/PlantEpigenome/index.html)

## Data Availability

scATAC-seq data encompassing 18 libraries from nine organs were accessible in NCBI (PRJNA1007577/GSE252040; https://dataview.ncbi.nlm.nih.gov/object/PRJNA1007577?reviewer=kgarq48dii11vomg44kgr1jq66; PRJNA1052039; https://dataview.ncbi.nlm.nih.gov/object/PRJNA1052039?reviewer=flhu9sl84o5m999r1ph8tlmmbg)

## Acknowledgements

This research was funded by the National Science Foundation (IOS-2134912) to SRW and RJS and the UGA Office of Research to RJS. APM and JPM were supported by the National Institutes of Health (K99GM144742) and (T32GM142623), respectively. DW was supported by National Science Foundation MCB-2224729 and IOS-1953279. We thank Changhui Sun (Rice Research Institute, Sichuan Agricultural University, Chengdu 611130, China) for identifying genetic variants of ZS97 using NIP as reference genotype in *O. sativa*.

## Contributions

R.J.S., H.Y., J.P.M., A.P.M., M.A.A.M., D.W., S.Z., and S.R.W. designed and conceived experiments and managed the project. X.Z., Y.L., Z.L., X.T., S.Z., Y.W., and H.Y. participated in material collection and sample processing. H.Y., J.P.M., and T.R. performed the bioinformatics analyses. H.Y. and J.P.M. wrote the manuscript. Y.L. Y.W. and Z.L. contributed to marker validation. R.J.S., J.P.M., A.P.M., M.A.A.M., D.W., S.Z., and S.R.W. edited the manuscript. L.H. contributed to some computing resources.

## Supplementary information

### Supplementary Note

1. Cell-type annotation and validation

### Supplementary Tables

Supplementary Tables 1–27.

---

**Extended Data Fig. 1.**
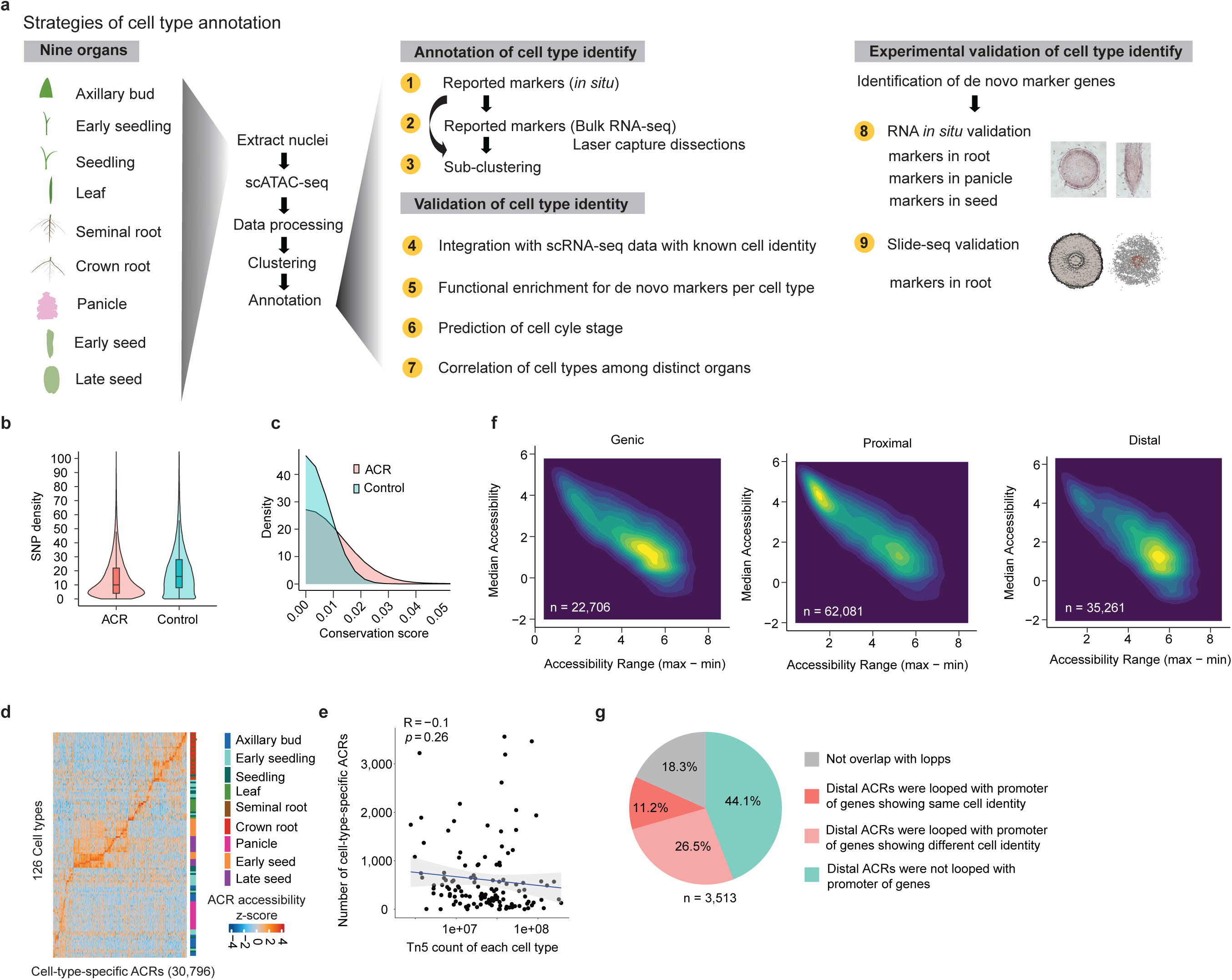
Pipeline of cell identity annotation and quality control of rice ACRs. **a,** A pipeline of cell identity annotation corresponding to nine strategies shown in Supplementary Note 1. **b-c,** A comparison between ACRs (n = 120,048) and control regions (n = 120,048) in terms of SNP density (**b**) and conservation scores (**c**). The SNP data were downloaded from Rice SNP-Seek Database (Mansueto et al. 2017). The SNP density is the number of SNPs per 1,000 bp within the ACRs or control regions. The ACRs exhibited lower SNP density and higher conservation scores compared to a control composed of non-ACR DNA. The control set was created to match each ACR by excluding exons, ACRs, and unmappable regions, and they possess a similar GC content (< 5% average difference) compared to the positive set. **d,** Row normalized chromatin accessibility z-scores for ACRs across cell types. **e,** Spearman correlation is shown between the total number of Tn5 insertions and number of cell-type-specific ACRs across all cell types. The identification of cell-type-specific ACRs was independent of sequencing depth, as evidenced by the lack of correlation between Tn5 insertions and the number of cell-type-specific ACRs per cell type. **f,** A two-dimensional density plot illustrates the median chromatin accessibility across 126 cell types for 120,048 ACRs, along with the range of chromatin accessibility (calculated as the difference between the maximum and minimum values). **g,** Categories of distal cell-type-specific leaf ACRs showing potential interaction with genes based on examining chromatin loops derived from rice leaf Hi-C data.

**Extended Data Fig. 2.**
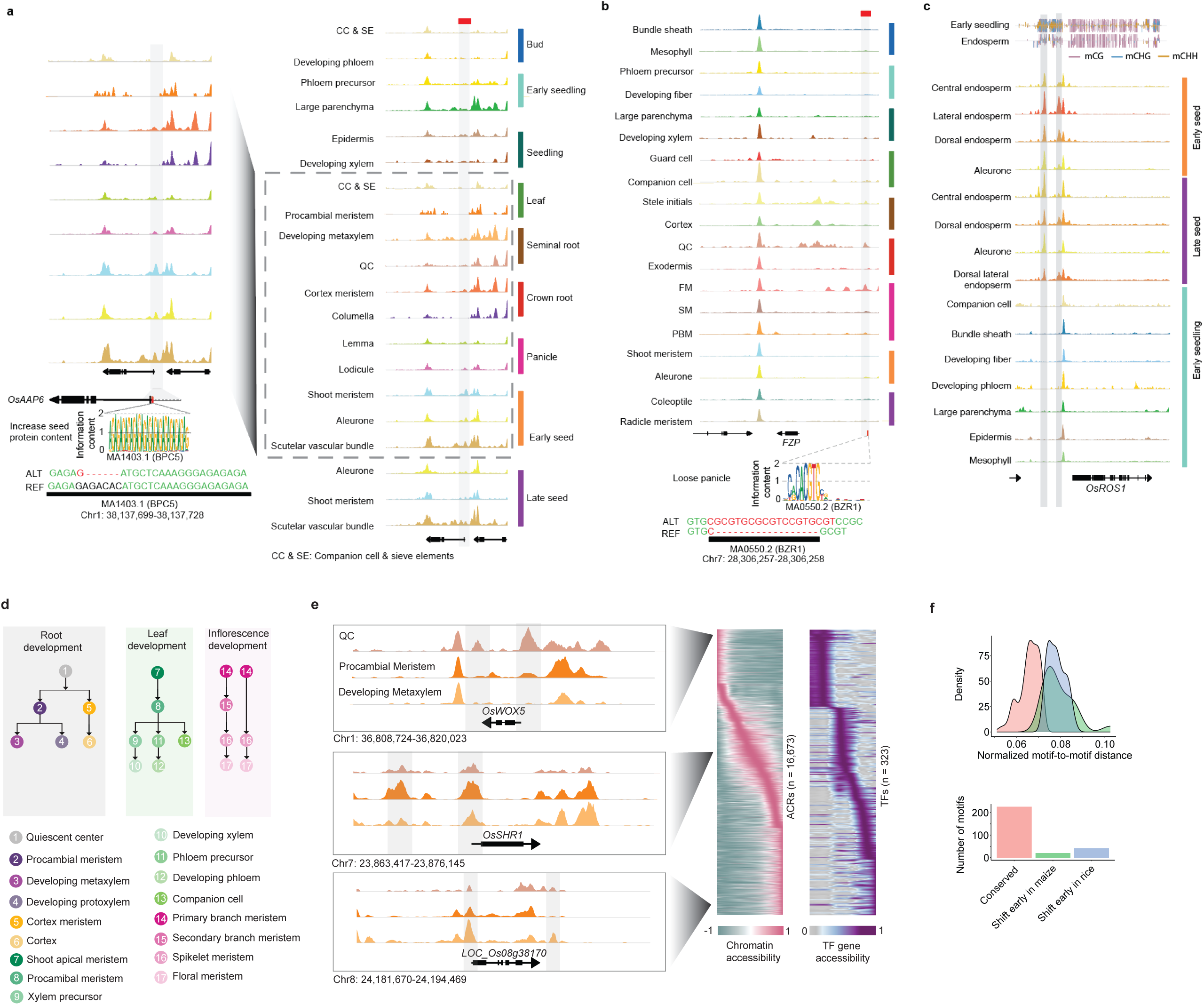
Characterization of QTNs located within cell-type-specific ACRs and pseudotime trajectories. **a,** The ratio of non-CDS QTNs overlapping with ACRs to all non-CDS QTNs. **b,** A bar plot reveals the non-CDS QTNs significantly overlapped ACRs. **c,** Analysis of cell-type-aggregate chromatin accessibility of ACRs surrounding *OsROS1* across endosperm-related cell types and cell types in early seedling tissue. The gray bar highlights endosperm-specific cytosine methylation depletion over endosperm-specific ACRs. **d,** Overview of pseudotime developmental trajectory analysis. **e,** Relative chromatin accessibility of ACRs and TF genes associated with pseudotime (x-axis). The left side displays three genes neighboring the enriched ACRs along the trajectory gradient. **f,** Distribution of motif-to-motif distances (upper panel) and number of motifs (lower panel) from k-mean groups.

**Extended Data Fig. 3.**
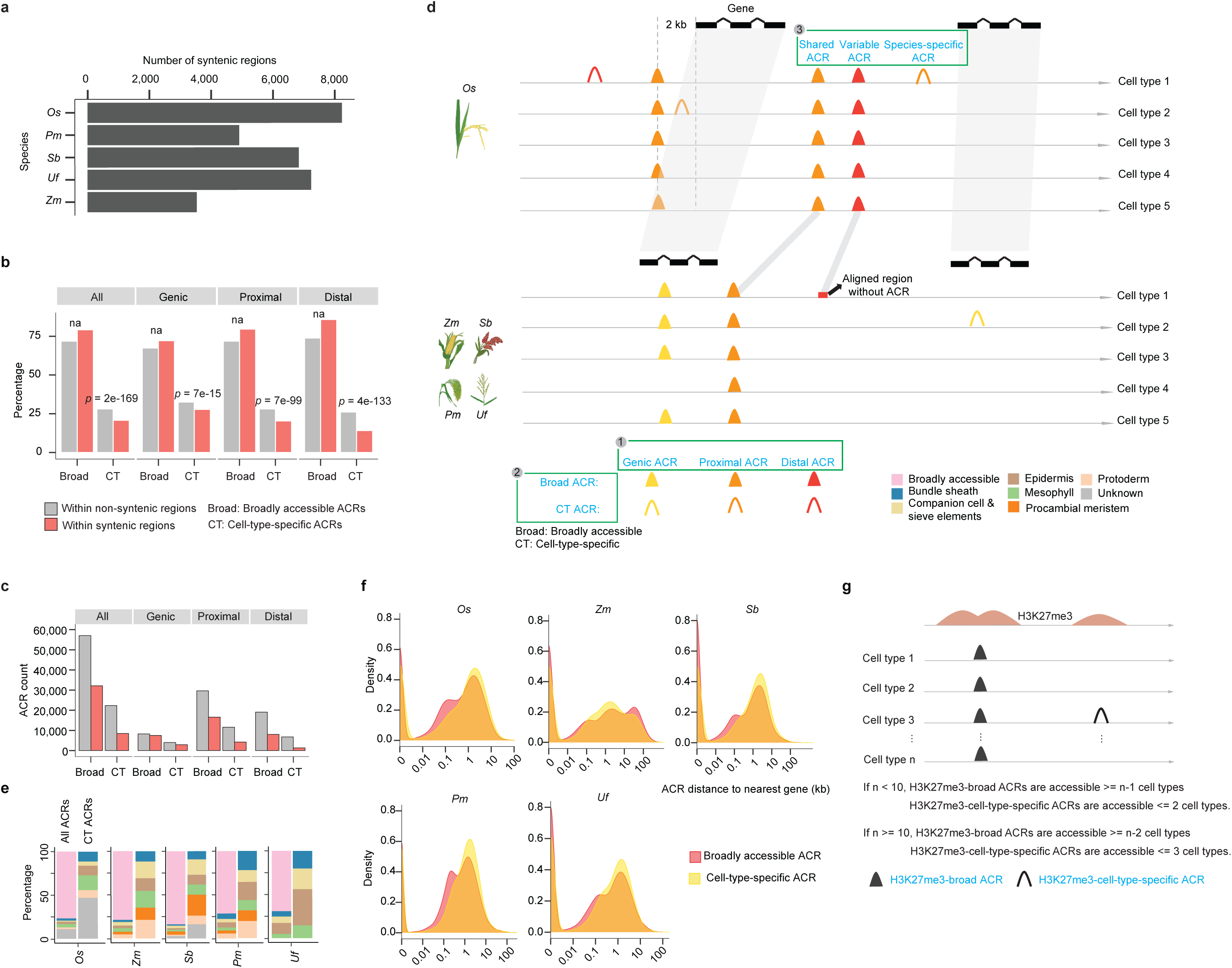
Characterization of ACRs across five grass species. **a,** Number of syntenic regions identified in all five species. Note that Os was used as the reference due to the inclusion of the atlas, biasing results to this segment of the tree. **b,** Enrichment of cell-type-specific ACRs in non-syntenic regions, compared to syntenic regions. Significance testing was performed using a two-sided Fisher’s exact test. ‘na’ indicates that the broadly-accessible ACRs were not enriched in non-syntenic regions for genic, proximal, and distal categories. The percentage for broad and cell-type-specific ACRs within non-syntenic or syntenic regions collectively sum to 100%. **c,** ACR count of broadly-accessible and cell-type-specific ACRs within syntenic and non-syntenic regions. **d,** A summary plot displaying the different categories of ACRs: 1) Based on their proximity to the nearest gene, ACRs are divided into three groups: genic ACRs (overlapping a gene), proximal ACRs (within 2 kb of genes), and distal ACRs (more than 2 kb away from genes). 2) According to the statistical approach described in the methods section (Identification of cell-type-specific ACRs in leaf tissue across five species), ACRs are classified as either broad ACRs, which are accessible in most cell types (*p* value >= 0.0001), or cell-type-specific ACRs, which are accessible in a limited number of cell types (*p* value < 0.0001). 3) Based on their syntenic status, ACRs are grouped into shared ACRs (matching sequences accessible in both species), variable ACRs (matching sequences accessible in only one species), and species-specific ACRs (sequences exclusive to a single species). **e,** The distribution of cell-type-specific ACRs across different cell types was characterized by a similar percentage representation. **f,** The distribution of distance of ACRs to their nearest genes in broadly accessible and cell-type-specific ACRs across five species. **g,** A summary plot displaying criteria to define H3K27me3-broad and H3K27me3-cell-type-specific ACRs. For the analysis of cell types in leaf tissue across rice, sorghum, and maize, where the number of cell types analyzed is fewer than 10, we set criteria such that H3K27me3-broad ACRs must be accessible in at least n-1 cell types, whereas cell-type-specific ACRs should be accessible in fewer than 3 out of the n examined cell types. For seedling and seminal root tissues in rice, where more than 10 cell types are examined, we adjusted the criteria. Here, H3K27me3-broad ACRs are required to be accessible in at least n-2 cell types, and cell-type-specific ACRs should be accessible in fewer than 4 out of the n examined cell types.

**Extended Data Fig. 4.**
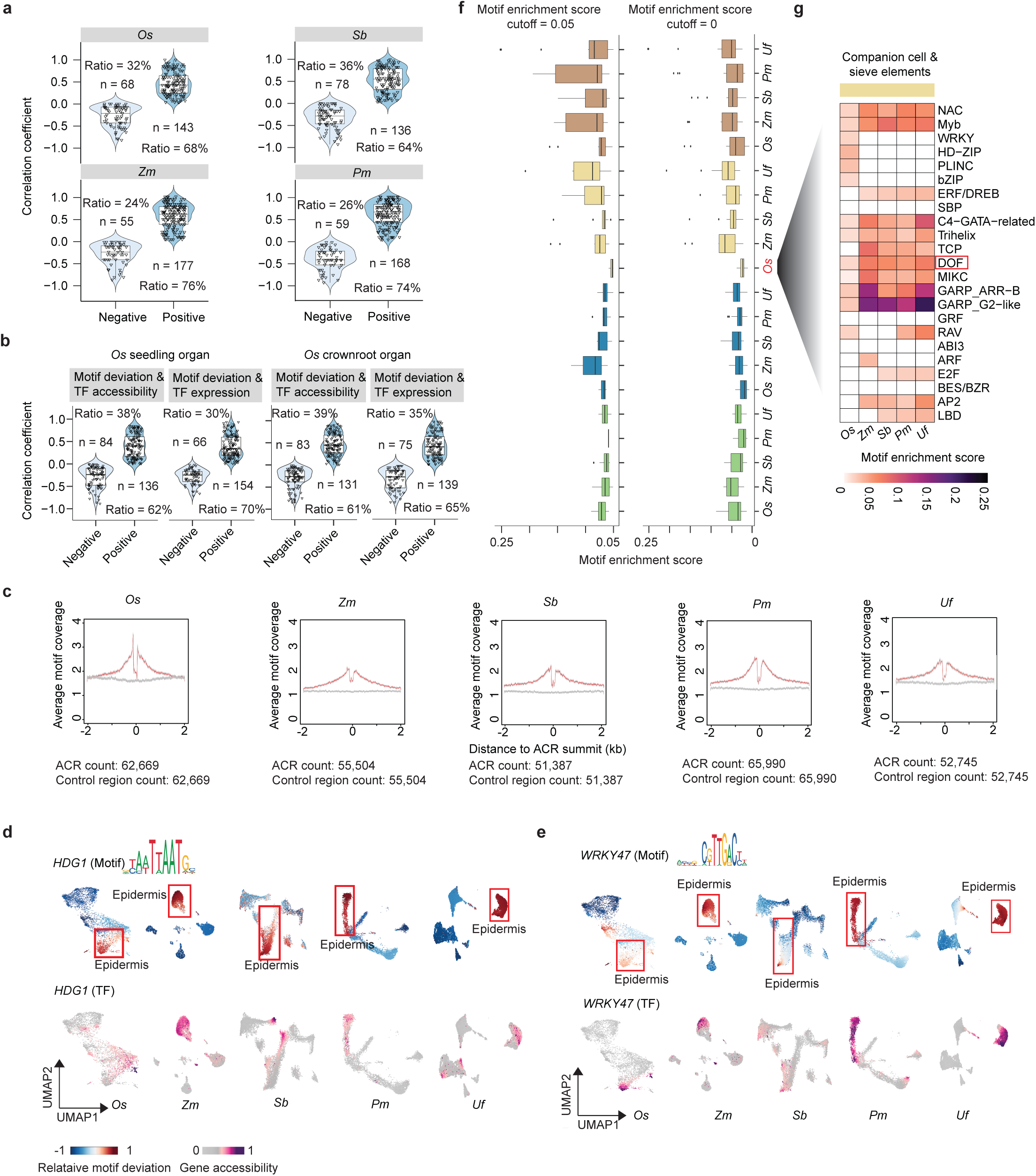
Examination of the chromatin accessibility of motifs enriched in specific cell types across species. **a,** Spearman correlation between chromatin accessibility of TF genes and motif deviation. Note that for *U. fusca* species, the analysis was omitted due to the limited number of cell types (four) available for spearman correlation, making it statistically unfeasible. **b,** Spearman correlation of motif deviation to chromatin accessibility and expression of TF genes in *O. sativa* seedling organ. **c,** Mean motif coverage centered on ACRs (red line) and control regions (gray line) for each species, with 95% confidence intervals shown as a shaded polygon. **d,** The UMAP panels highlight one TF motif (HDG1) enriched with ACRs in epidermis, where their cognate TFs are known to accumulate. **e,** The UMAP panels highlight one representative TF motif underlying the WRKY family enriched with ACRs in epidermis cells, where their cognate TFs are known to accumulate. **f-g,** Presenting box (**e**) and heatmap (**f**) plots illustrating the collapsed motif enrichment patterns into TF motif families across various species for each cell type. Each dot within the box represents a TF motif family. The score for each super TF motif family was calculated by averaging the enrichment scores of all the TF motif members within that super family. The DOF TF motif family was highlighted by a red frame in the heatmap. The reduced motif enrichment cutoff setting reveals additional motif families enriched in companion cells and sieve elements, compared to the higher beta cutoff setting as shown in Fig. 3e. It is worth noting that we still observed lower enrichment scores in *O. sativa* compared to other species.

**Extended Data Fig. 5.**
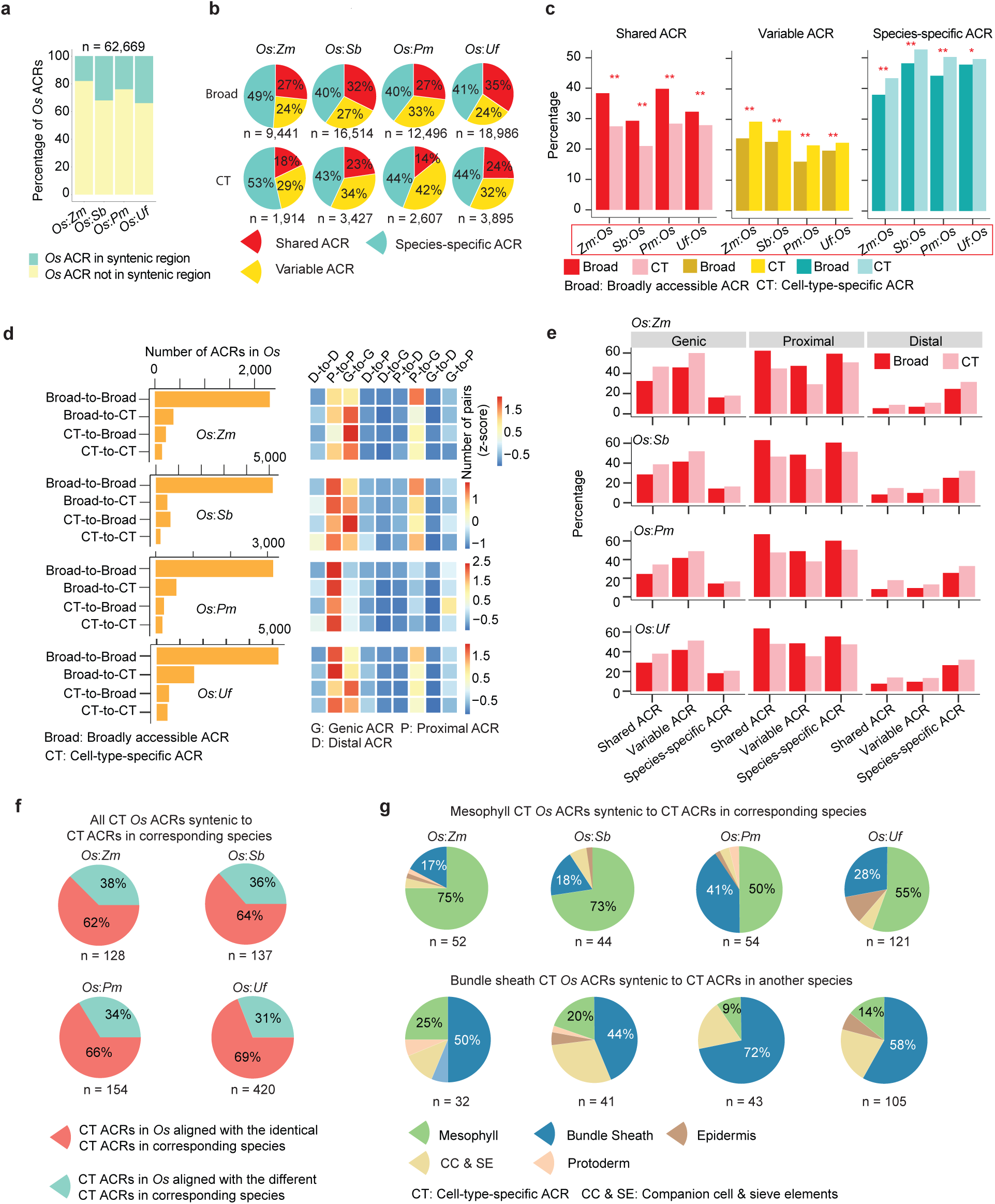
Characterization of ACRs within syntenic regions. **a,** The bar plot illustrates the percentage of *O. sativa* ACRs within syntenic and non-syntenic regions when compared to another species. **b,** The pie charts depict percentages of broadly accessible (Broad) and cell-type-specific (CT) ACRs within three classes of ACR conservation in Fig. 3b. The total number of ACR in *O. sativa* is below the pie charts. **c,** The percentage of broad and cell-type-specific ACRs using *Z. mays*, *S. bicolor*, *P. miliaceum*, or *U. fusca* as the baseline underlying three classes shown in Fig. 3b. The x-axis labels, highlighted by a red frame, are the inverse to x-axis labels in Fig. 3c by using *O. sativa* as a baseline. The significance test was done by using the Fisher’s exact test (alternative = ‘two.sided’). ‘**’ denotes a *p* value < 0.01, and ‘*’ denotes a *p* value < 0.05. **d,** Characterization of ACRs under ‘shared ACR’ class and their proximity categorizations. **Left,** the number of ACRs under the ‘shared ACR’ class, categorized into four combinations based on broadly accessible and cell-type-specific ACRs. **Right,** a heatmap of ACR categories based on their proximity to genes: genic ACRs overlapping genes, proximal ACRs (within 2 kb of genes), and distal ACRs (more than 2 kb away from the TSS). **e,** The percentage of shared, variable, and species-specific ACR classes in genic, proximal, and distal manners for each species pair. The percentage for each class within genic, proximal, and distal groups collectively sum to 100%. E.g. Genic, proximal and distal broad shared ACRs sum to 100%. **f,** The pie charts illustrate percentage of cell-type-specific *O. sativa* ACRs syntenic to cell-type-specific ACRs in corresponding species. **g,** The pie charts illustrate mesophyll and bundle sheath specific *O. sativa* ACRs syntenic to cell-type-specific ACRs in corresponding species.

**Extended Data Fig. 6.**
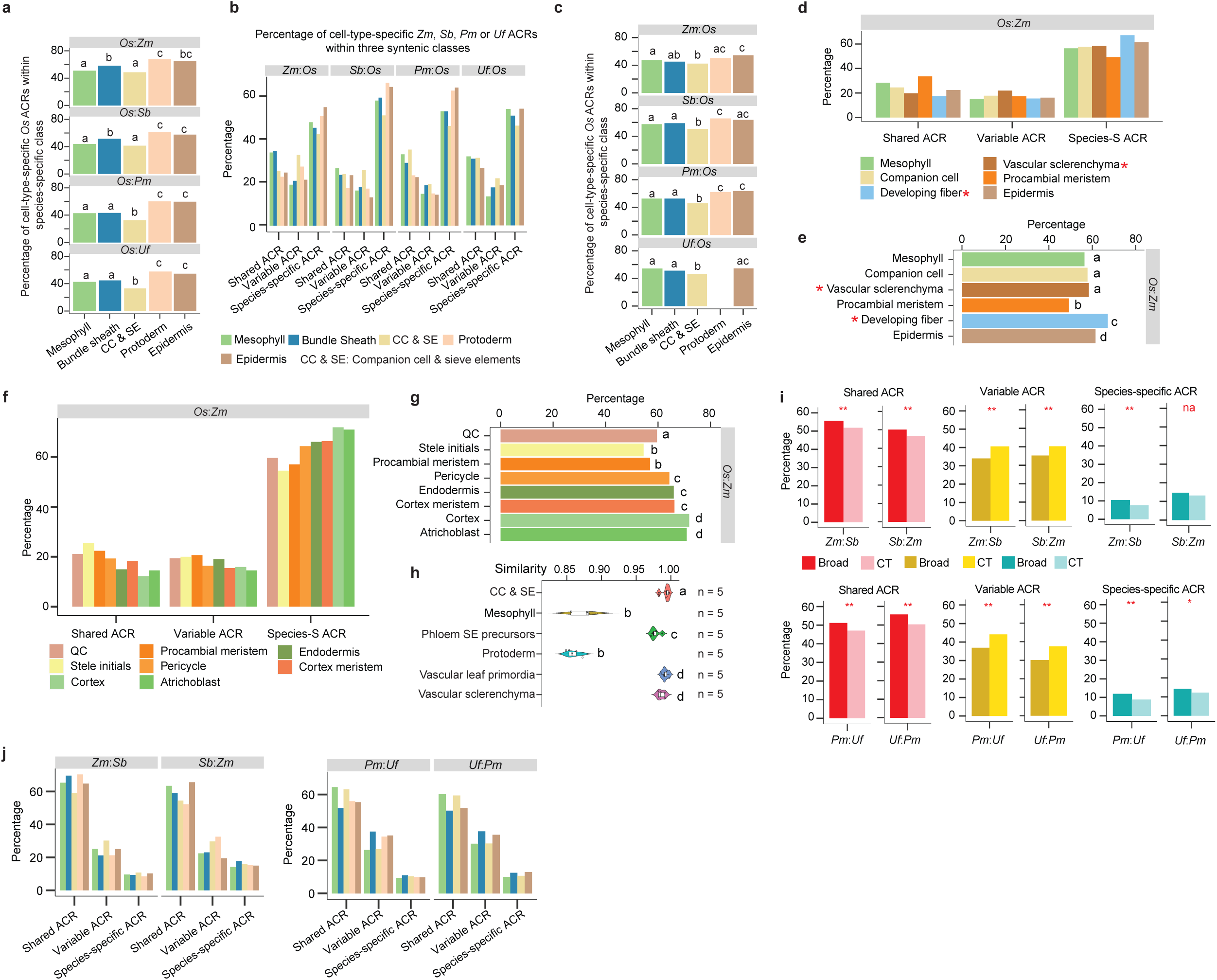
Epidermal cell specific ACRs are less conserved in sequence than other examined cell types. **a,** The analysis demonstrates an enrichment of cell-type-specific ACRs based on Fisher’s exact test. This test assesses whether cell-type-specific ACRs are more likely to be situated in the ‘species-specific’ class, compared to other cell types. Statistically significant differences (*p* value < 0.05) in all pairwise comparisons are denoted by distinct letters, determined using Fisher’s exact test with the alternative set to ‘two.sided’. Bars sharing the same letter indicate that they are not significantly (*p* value > 0.05) different from each other. **b,** The proportion of cell-type-specific ACRs identified collectively across all cell types within each species pair under three syntenic classes. The percentage for each cell type within the three classes collectively sum to 100%. **c,** This panel has the same meaning as panel **a**, and corresponds to an enrichment test on panel **b**. *U. fusca*-to-*O. sativa* did not include a bar for protoderm as this cell type has not been identified in *U. fusca* scATAC-seq dataset. **d,** The proportion of cell-type-specific ACRs identified collectively across all cell types within each species pair under three syntenic classes (share, variable, and species-specific classes). The percentage for each cell type within the three classes collectively sum to 100%. The asterisk indicates three cell types not examined in the comparisons within leaf tissue presented in Fig. 4e. **e,** This test assesses whether cell-type-specific ACRs are more likely to be situated in the ‘species-specific’ class, compared to other cell types. Statistically significant differences (*p* value < 0.05) in all pairwise comparisons are denoted by distinct letters, determined using a Fisher’s exact test with the alternative set to ‘two.sided’. Bars sharing the same letter indicate that they are not significantly (*p* value > 0.05) different from each other. The asterisk indicates three cell types not examined in the comparisons within leaf tissue presented in Fig. 4e. **f,** The proportion of cell-type-specific ACRs identified collectively across all cell types within each species pair under three syntenic classes (share, variable, and species-specific classes). The percentage for each cell type within the three classes collectively sum to 100%. **g,** This test assesses whether cell-type-specific ACRs are more likely to be situated in the ‘species-specific’ class, compared to other cell types. Statistically significant differences (*p* value < 0.05) in all pairwise comparisons are denoted by distinct letters, determined using Fisher’s exact test with the alternative set to ‘two.sided’. Bars sharing the same letter indicate that they are not significantly (*p* value > 0.05) different from each other. **h,** The MetaNeighbor analysis quantifies transcriptome similarity among cell types in *O. sativa* compared to *Z. mays*. ‘n=5’ denotes five replicates, which were created by randomly dividing the cells from each cell type into five groups, with each group serving as input for the MetaNeighbor analysis. Statistically significant differences (*p* value < 0.05) in all pairwise comparisons are denoted by distinct letters, determined using Wilcoxon signed rank test with the alternative set to ‘two.sided’. The groups sharing the same letter indicate that they are not significantly (*p* value > 0.05) different from each other. **i,** The percentage of broad and cell-type-specific ACRs using either *Z. mays* (*Zm*) or *S. bicolor* (*Sb*) as the baseline and using either *P. miliaceum* (*Pm*) or *U. fusca* (*Uf*) as the baseline underlying three classes shown in Fig. 4b. The significance test was done by using the Fisher’s exact test (alternative = ‘two.sided’). ‘**’ denotes a p value < 0.01, and ‘*’ denotes a *p* value < 0.05. **j,** The percentage of cell-type-specific ACRs identified across all cell types within each species pair split into three classes shown in Fig. 4b. The percentage for each cell type within the three classes collectively sum to 100%.

**Extended Data Fig. 7.**
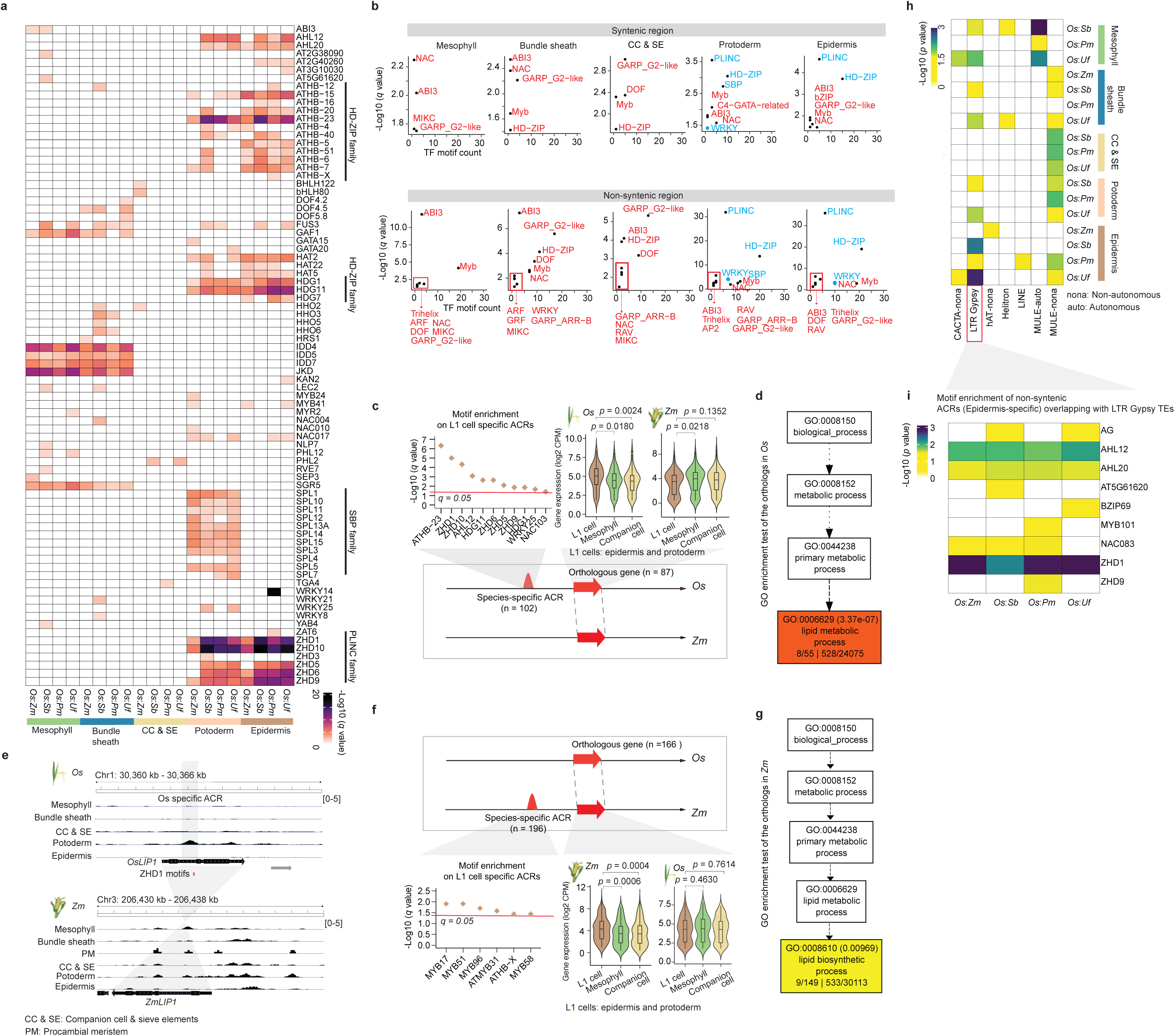
Identification of TF motifs and TEs associating with epidermal cell divergence. **a,** TF motif enrichment tests were performed in species-specific ACRs per species pair using *O. sativa* as the baseline (See Methods: Binomial test-based motif enrichment analysis). The TF motif with a *q* value more than 0.05 was indicated by filling them with a white color. **b,** This panel utilized the same method as panel a to conduct TF motif enrichment tests in syntenic and non-syntenic ACRs in the *O. sativa*-to-*Z. mays* species pair (The results from other species pairs were shown in Supplementary Table 8). The ‘-Log10 (*q* value)’ of each TF family was calculated by averaging ‘-Log10 (*q* value)’ across all individual TF motifs within this family. TF family names were marked besides each dark dot, while the PLINC, HD-ZIP, SBP, and WRKY family were highlighted using blue color. The x-axis indicates the number of TF motifs detected to be enriched (*q* value < 0.05) in the relative TF family. **c,** TF motif enrichment tests were performed in species-specific ACRs neighboring *O. sativa* ortholog exhibiting higher expression levels in epidermis based on Binomial test (See Methods: Binomial test-based motif enrichment analysis). Significance testing in violin plot was performed using the t-test (alternative = ‘two.sided’). **d,** GO enrichment test was performed in *O. sativa* orthologous genes based on agriGO (Tian et al. 2017). **e,** A screenshot of *LIP1* accessibility in *O. sativa* and *Z. mays* L1 cells which contains an *O. sativa* epidermal specific and species-specific ACR with two ZHD1 motif sites. No corresponding ZHD1 motifs were found in *ZmLIP1*. **f-g,** These two plots have the same meanings as panel **d** and **e**, but focusing on species-specific ACRs in *Z. mays*. **h,** Enrichment of TE family on ACRs within non-syntenic regions relative to those in syntenic regions. Significance testing was performed using Fisher’s exact test (alternative = ‘two.sided’). The TE with a *p* value more than 0.05 was indicated by filling them with a white color. **i,** TF motif enrichment tests were performed on the epidermis specific ACRs overlapping with LTR *Gypsy* TEs based on Binomial test (See Methods: Binomial test-based motif enrichment analysis). The TE with a *p* value more than 0.05 was indicated by filling them with a white color.

**Extended Data Fig. 8.**
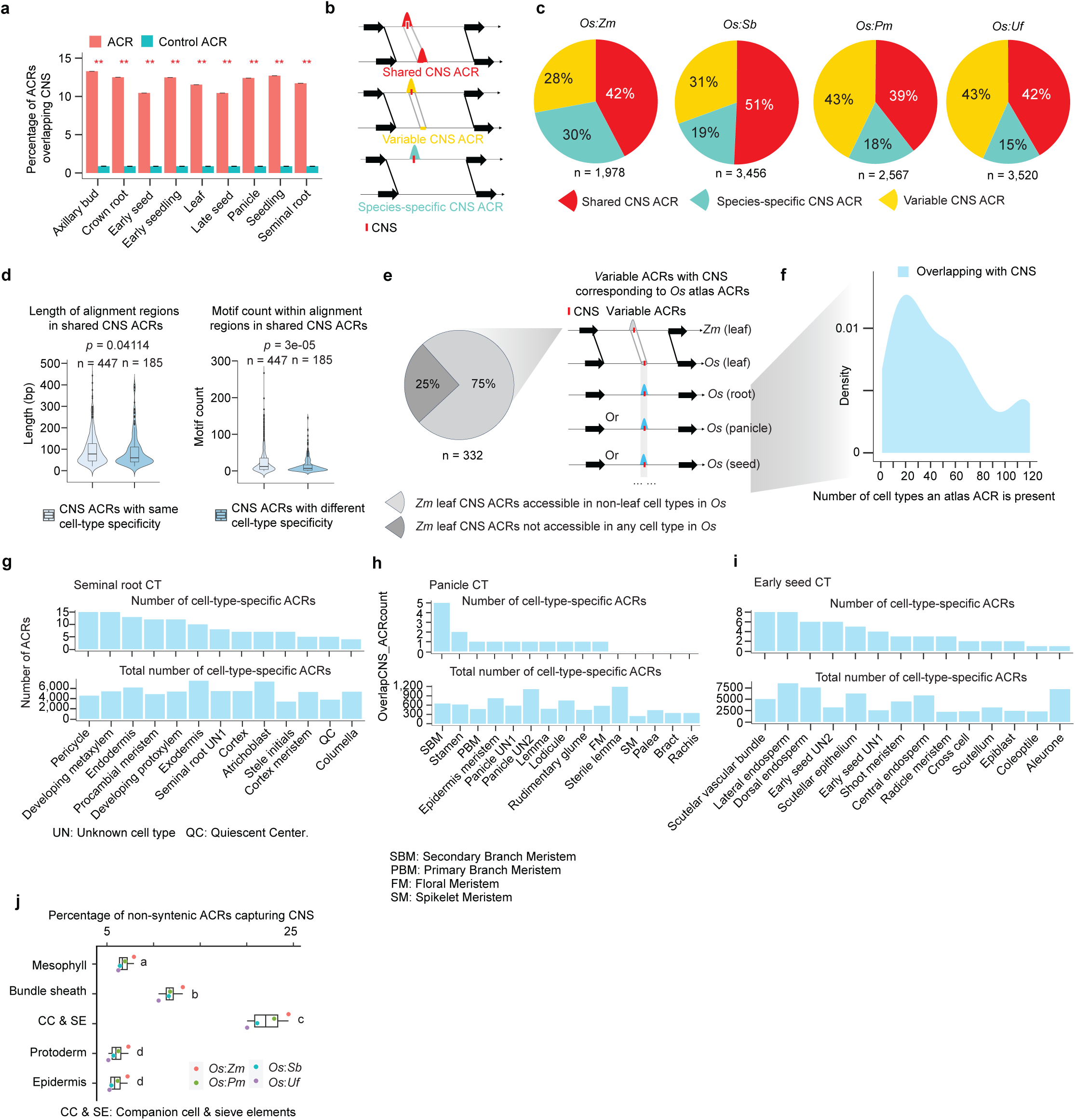
Examination of *O. sativa* atlas ACRs corresponding to variable *Z. mays* ACRs. **a,** Percentage of ACRs overlapping CNS. The significance test was done by using the Binomial test (** indicate p value < 0.01; alternative = ‘two.sided’). We generated control sets by simulating sequences with the same length as ACRs 100 times, yielding a mean proportion for the control sets. The binomial test p value was calculated by comparing the mean ratio to the observed overlapping ratio of ACRs capturing the CNS. **b,** Three classes depicting variations in ACR conservation between two species. ‘Shared CNS ACRs’: CNS ACRs with matching sequences that are accessible in both species; ‘Variable CNS ACRs’: CNS ACRs with matching sequences, but are only accessible in one species; ‘Species-specific CNS ACRs’: CNS ACRs where the sequence is exclusive to a single species. **c,** The pie charts depict percentages of CNS ACRs within three classes of ACR conservation in panel **a**. The total number of ACR in *O. sativa* is below the pie charts. **d,** Comparison of length of alignment regions and motif count within shared CNS ACRs between with same and different cell-type specificity **e,** A sketch illustrating whether variable ACRs containing CNS in *Z. mays* capture ACRs derived from the *O. sativa* atlas. The left pie chart panel represents the percentage of *Z. mays* leaf CNS ACRs that were accessible in non-leaf cell types in the *O. sativa* atlas. **f,** A density plot illustrating the number of non-leaf *O. sativa* cell-types in which an *O. sativa* ACR syntenic to *Z. mays* variable ACRs are accessible. **g-i,** The bar plots represent the count of *O. sativa* atlas cell-type-specific ACRs accessible in non-leaf cell states from seminal root, panicle, and early seed organs. These ACRs were derived from cell-type-specific ACRs from Fig. 4e. **j,** The percentage of non-syntenic ACRs capturing CNS. All pairwise comparisons are statistically significant (*p* value < 0.05) as indicated by different letters based on t-test (alternative = ‘two.sided’). If two bars have the same letter, then they are equivalent (*p* value > 0.05).

**Extended Data Fig. 9.**
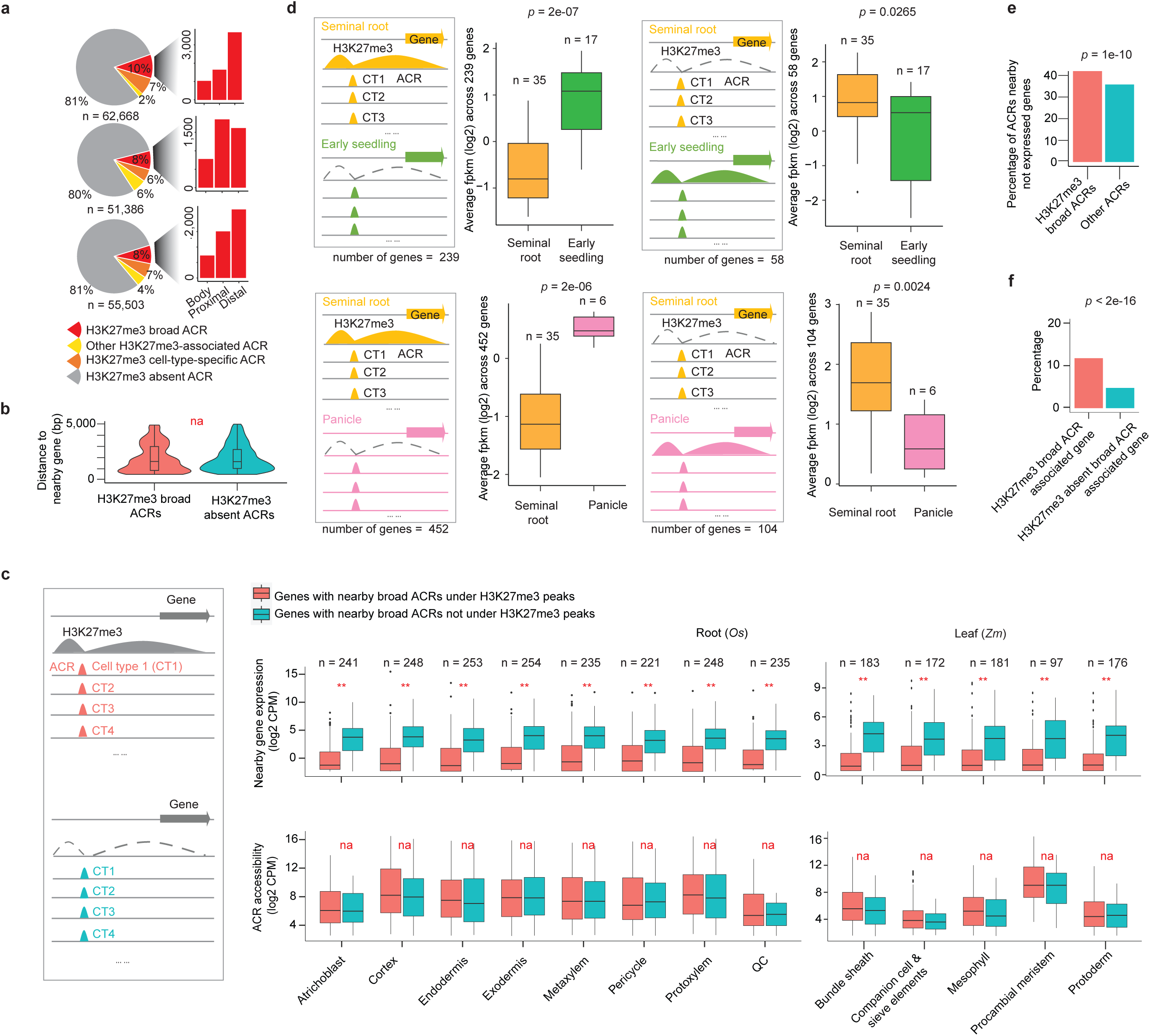
Assessment of transcriptional state of genes in proximity to H3K27me3-broad ACRs. **a,** Pie charts showcasing the composition of different categories of ACRs, and bar plots displaying the number of ACRs within the H3K27me3-broad-ACR group. **b,** No significant difference in the distance to nearby genes was identified between H3K27me3-broad ACRs to H3K27me3-absent ACRs as shown in Fig. 5c. **c,** Comparative analysis of expression levels and chromatin accessibility of genes surrounding broad ACRs under and outside of H3K27me3 peaks. ** indicate *p* value < 0.01, which was performed by the Wilcoxon signed rank test (alternative = ‘two.sided’). **d,** Four categories of gene sets overlapped with H3K27me3 peaks and comparisons of their expression between bulk RNA-seq data libraries from two different organs. The ‘n’ above the box indicates the number of libraries used in this comparison. Significance testing was performed using the Wilcoxon signed rank test (alternative = ‘two.sided’). **e,** Percentage of H3K27me3-broad ACRs nearby genes exhibiting no expression across any cell type was significantly higher than the other ACRs associated with the not expressed genes. Significance testing was performed using Fisher’s exact test (alternative = ‘two.sided’). **f,** Percentage of H3K27me3-broad ACRs nearby genes exhibiting cell-type-specific expression was significantly higher than the other genes not associating with H3K27me3-broad ACRs. Significance testing was performed using Fisher’s exact test (alternative = ‘two.sided’).

**Extended Data Fig. 10.**
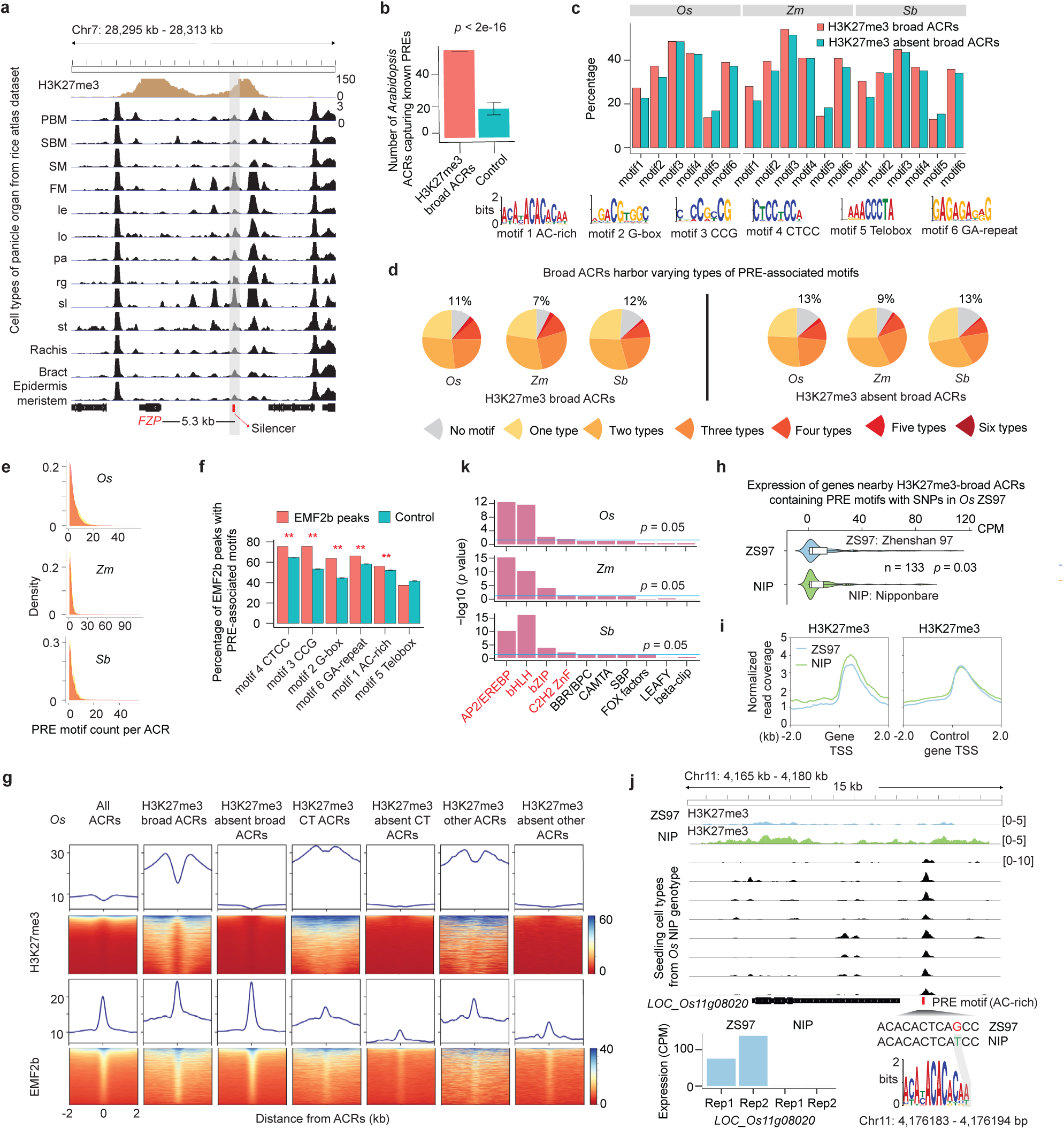
Characterization of H3K27me3-broad ACRs potentially enriched for silencer CREs. **a,** A screenshot illustrates a H3K27me3-broad ACR harboring a reported silencer within an H3K27me3 peak in the panicle organ, located approximately 5.3 kb upstream of *FZP* gene. SBM: Secondary Branch Meristem. PBM: Primary Branch Meristem. SM: Spikelet Meristem. FM: Floral Meristem. le: Lemma. lo: Lodicule. pa: Palea. rg: Rudimentary glume. sl: Sterile lemma. **b,** The number of H3K27me3-broad-ACR capturing known 53 PREs is determined based on *A. thaliana* scATAC-seq data. The significance test was done by using the Binomial test (alternative = ‘two.sided’; See Methods: Construction of control sets for enrichment tests). **c,** Percentage of H3K27me3 and H3K27me3 absent in broad ACRs that contain PRE-associated motifs. **d,** Pie charts illustrate the different types of PRE-associated motifs within H3K27me3-broad and H3K27me3-absent-broad ACRs. **e,** Distribution of PRE motif count per ACR for H3K27me3-broad and H3K27me3-absent-broad ACRs categories. **f,** Percentage of EMF2b peaks in *O. sativa* capturing six known motifs enriched in PREs in *A. thaliana*. ** indicate *p* value < 0.01, which was performed by the Binomial test (alternative = ‘two.sided’; See Methods: Construction of control sets for enrichment tests). **g,** Alignment of H3K27me3 and EMF2b chromatin attributes at summits of distinct ACR groups in *O. sativa*. CT ACRs mean cell-type-specific ACRs, other ACR means ACRs not classified as the broad or CT ACRs according to the classification criteria in Extended Data Fig. 3g. **h,** Comparison of expression of genes nearby H3K27me3-broad ACRs that contain PRE motifs with SNPs in ‘Zhenshan 97’ (ZS97) genotype using ‘Nipponbare’ (NIP) as the reference. Significant test was performed by the Wilcoxon signed rank test (alternative = ‘two.sided’). CPM: Counts Per Million. **i,** Alignment of H3K27me3 chromatin attributes at summits of transcription start site (TSS) of genes derived from panel h. The control genes include genes overlapping with H3K27me3, which are shared between both genotypes. **j,** A screenshot illustrates an H3K27me3-broad ACR containing a PRE-associated motif with an SNP situated at 1.2 kb upstream of *LOC_Os11g08020* gene, which might be associated with a lower H3K27me3 signal and higher expression of the *LOC_Os11g08020* in the ‘Zhenshan 97’ genotype. **k,** Four TF families, highlighted in red, were significantly enriched in H3K27me3-broad ACRs. The motif data were collected from 568 TFs from *A. thaliana* belonging to 24 families within the JASPAR database (Castro-Mondragon et al. 2022). The *p* value was computed using a hypergeometric test (alternative = ‘two.sided’).

**Extended Data Fig. 11.**
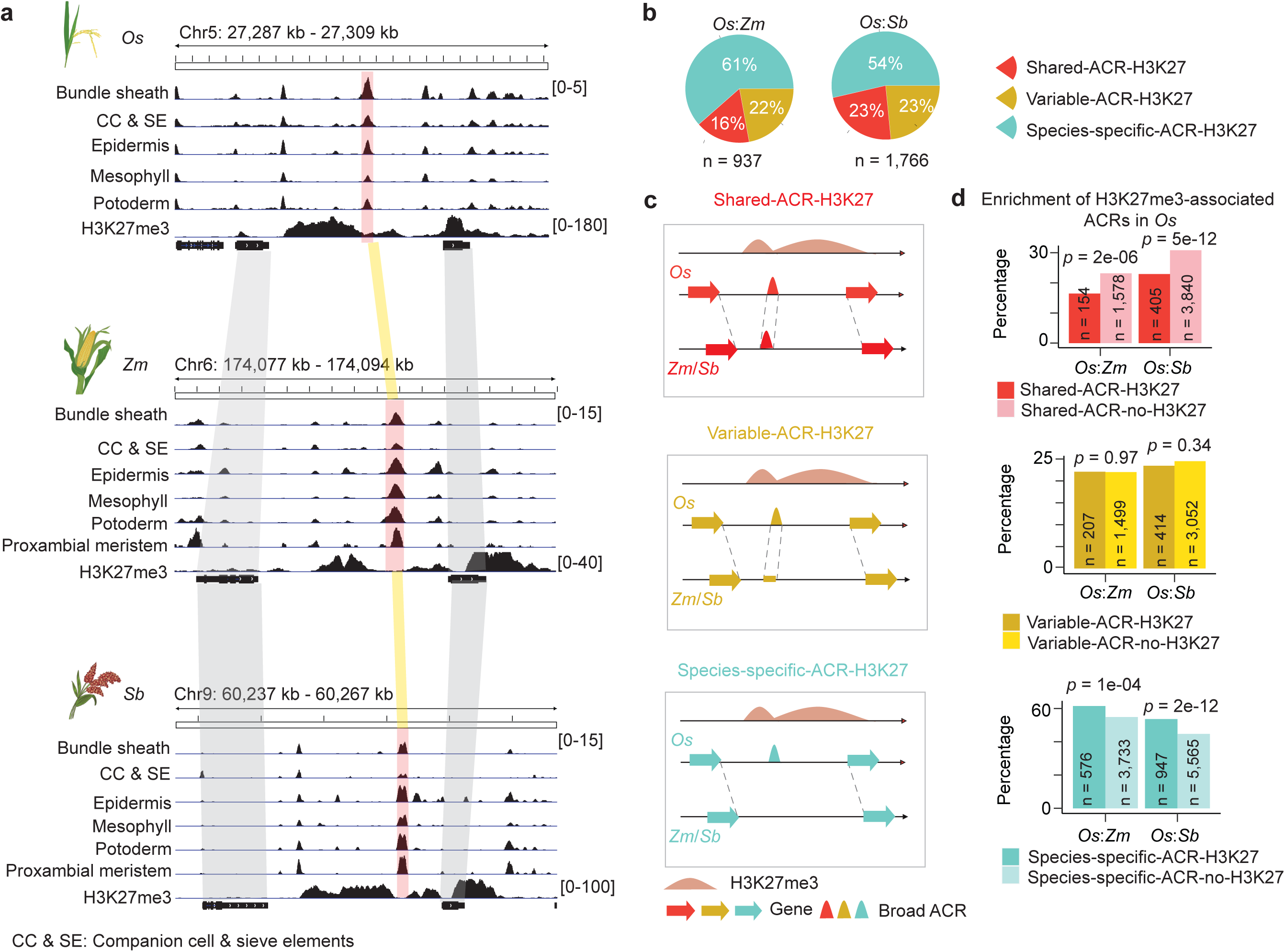
Evolutionary dynamics of H3K27me3-broad ACRs across different species. **a,** An example of a syntenic region containing H3K27me3-broad ACRs that were conserved across *O. sativa*, *Z. mays*, and *S. bicolor*. **b,** The pie charts depict percentages of H3K27me3-broad ACRs within three classes of ACR conservation in Fig. 3b. **c,** Three categories of *O. sativa* H3K27me3-broad ACRs that are syntenic to regions from another species. **d,** Comparison of ACR percentage underlying three classes corresponding to panel **c** associated with H3K27me3 or not. The *p* values displayed in the bottom bar plot were computed using Fisher’s exact test (alternative = ‘two.sided’).

## References

1 Preissl, S., Gaulton, K. J. & Ren, B. Characterizing cis-regulatory elements using single-cell epigenomics. Nature Reviews Genetics 24, 21–43 (2023).

2 Wittkopp, P. J. & Kalay, G. Cis-regulatory elements: molecular mechanisms and evolutionary processes underlying divergence. Nature Reviews Genetics 13, 59–69 (2012).

3 Marand, A. P., Eveland, A. L., Kaufmann, K. & Springer, N. M. cis-Regulatory elements in plant development, adaptation, and evolution. Annual review of plant biology 74, 111–137 (2023).

4 Oka, R. et al. Genome-wide mapping of transcriptional enhancer candidates using DNA and chromatin features in maize. Genome biology 18, 1–24 (2017).

5 Maher, K. A. et al. Profiling of accessible chromatin regions across multiple plant species and cell types reveals common gene regulatory principles and new control modules. The Plant Cell 30, 15–36 (2018).

6 Lu, Z. et al. The prevalence, evolution and chromatin signatures of plant regulatory elements. Nature Plants 5, 1250–1259 (2019).

7 Reynoso, M. A. et al. Evolutionary flexibility in flooding response circuitry in angiosperms. Science 365, 1291–1295 (2019).

8 Kajala, K. et al. Innovation, conservation, and repurposing of gene function in root cell type development. Cell 184, 3333–3348. e3319 (2021).

9 Ricci, W. A. et al. Widespread long-range cis-regulatory elements in the maize genome. Nature plants 5, 1237-1249 (2019).

10 Cusanovich, D. A. et al. A single-cell atlas of in vivo mammalian chromatin accessibility. Cell 174, 1309–1324. e1318 (2018).

11 Domcke, S. et al. A human cell atlas of fetal chromatin accessibility. Science 370, eaba7612 (2020).

12 Lu, Z. et al. Tracking cell-type-specific temporal dynamics in human and mouse brains. Cell 186, 4345–4364. e4324 (2023).

13 Javelle, M., Vernoud, V., Rogowsky, P. M. & Ingram, G. C. Epidermis: the formation and functions of a fundamental plant tissue. New Phytologist 189, 17–39 (2011).

14 Kadioglu, A., Terzi, R., Saruhan, N. & Saglam, A. Current advances in the investigation of leaf rolling caused by biotic and abiotic stress factors. Plant Science 182, 42–48 (2012).

15 Xu, Y. et al. Overexpression of OsZHD1, a zinc finger homeodomain class homeobox transcription factor, induces abaxially curled and drooping leaf in rice. Planta 239, 803–816 (2014).

16 Dorrity, M. W. et al. The regulatory landscape of Arabidopsis thaliana roots at single-cell resolution. Nature communications 12, 3334 (2021).

17 Farmer, A., Thibivilliers, S., Ryu, K. H., Schiefelbein, J. & Libault, M. Single-nucleus RNA and ATAC sequencing reveals the impact of chromatin accessibility on gene expression in Arabidopsis roots at the single-cell level. Molecular Plant 14, 372–383 (2021).

18 Tu, X., Marand, A. P., Schmitz, R. J. & Zhong, S. A combinatorial indexing strategy for low-cost epigenomic profiling of plant single cells. Plant Communications 3 (2022).

19 Nobori, T. et al. Time-resolved single-cell and spatial gene regulatory atlas of plants under pathogen attack. bioRxiv, 2023.2004. 2010.536170 (2023).

20 Marand, A. P., Chen, Z., Gallavotti, A. & Schmitz, R. J. A cis-regulatory atlas in maize at single-cell resolution. Cell 184, 3041–3055. e3021 (2021).

21 Feng, D. et al. Chromatin accessibility illuminates single-cell regulatory dynamics of rice root tips. BMC biology 20, 274 (2022).

22 Zhang, L. et al. Asymmetric gene expression and cell-type-specific regulatory networks in the root of bread wheat revealed by single-cell multiomics analysis. Genome Biology 24, 65 (2023).

23 Swift, J. et al. Single nuclei sequencing reveals C4 photosynthesis is based on rewiring of ancestral cell identity networks. bioRxiv, 2023.2010. 2026.562893 (2023).

24 Mendieta, J. P. et al. Investigating the cis-Regulatory Basis of C3 and C4 Photosynthesis in Grasses at Single-Cell Resolution. National Academy of Sciences of the United States of America 121, e2402781121 (2024).

25 Dong, Q. et al. Genome-wide Hi-C analysis reveals extensive hierarchical chromatin interactions in rice. The Plant Journal 94, 1141–1156 (2018).

26 Wei, X. et al. A quantitative genomics map of rice provides genetic insights and guides breeding. Nature Genetics 53, 243–253 (2021).

27 Yang, Y. et al. Natural variation of OsGluA2 is involved in grain protein content regulation in rice. Nature communications 10, 1949 (2019).

28 Zemach, A. et al. Local DNA hypomethylation activates genes in rice endosperm. Proceedings of the National Academy of Sciences 107, 18729–18734 (2010).

29 Xu, Q. et al. DNA demethylation affects imprinted gene expression in maize endosperm. Genome biology 23, 77 (2022).

30 Ohashi-Ito, K. & Fukuda, H. HD-Zip III homeobox genes that include a novel member, ZeHB-13 (Zinnia)/ATHB-15 (Arabidopsis), are involved in procambium and xylem cell differentiation. Plant and Cell Physiology 44, 1350-1358 (2003).

31 Wu, R. et al. CFL1, a WW domain protein, regulates cuticle development by modulating the function of HDG1, a class IV homeodomain transcription factor, in rice and Arabidopsis. The Plant Cell 23, 3392–3411 (2011).

32 Zhong, R., Richardson, E. A. & Ye, Z.-H. The MYB46 transcription factor is a direct target of SND1 and regulates secondary wall biosynthesis in Arabidopsis. The Plant Cell 19, 2776–2792 (2007).

33 Wang, Z. et al. Salicylic acid promotes quiescent center cell division through ROS accumulation and down-regulation of PLT1, PLT2, and WOX5. Journal of Integrative Plant Biology 63, 583-596 (2021).

34 Gontarek, B. C., Neelakandan, A. K., Wu, H. & Becraft, P. W. NKD transcription factors are central regulators of maize endosperm development. The Plant Cell 28, 2916–2936 (2016).

35 Sun, X. et al. Activation of secondary cell wall biosynthesis by miR319-targeted TCP 4 transcription factor. Plant Biotechnology Journal 15, 1284–1294 (2017).

36 Ortiz-Ramírez, C. et al. Ground tissue circuitry regulates organ complexity in maize and Setaria. Science 374, 1247–1252 (2021).

37 Lv, Z., Zhao, W., Kong, S., Li, L. & Lin, S. Overview of molecular mechanisms of plant leaf development: a systematic review. Frontiers in Plant Science 14 (2023).

38 Le Hir, R. & Bellini, C. The plant-specific Dof transcription factors family: new players involved in vascular system development and functioning in Arabidopsis. Frontiers in Plant Science 4, 164 (2013).

39 Dai, X. et al. Chromatin and regulatory differentiation between bundle sheath and mesophyll cells in maize. The Plant Journal 109, 675–692 (2022).

40 Yanagisawa, S. Dof1 and Dof2 transcription factors are associated with expression of multiple genes involved in carbon metabolism in maize. The Plant Journal 21, 281–288 (2000).

41 Yanagisawa, S. & Sheen, J. Involvement of maize Dof zinc finger proteins in tissue-specific and light-regulated gene expression. The Plant Cell 10, 75–89 (1998).

42 Borba, A. R. et al. Synergistic binding of bHLH transcription factors to the promoter of the maize NADP-ME gene used in C4 photosynthesis is based on an ancient code found in the ancestral C3 state. Molecular biology and evolution 35, 1690–1705 (2018).

43 Shimadzu, S., Furuya, T. & Kondo, Y. Molecular mechanisms underlying the establishment and maintenance of vascular stem cells in Arabidopsis thaliana. Plant and Cell Physiology 64, 274–283 (2023).

44 Otero, S. & Helariutta, Y. Companion cells: a diamond in the rough. Journal of experimental botany, erw392 (2016).

45 Ramachandran, V. et al. Plant-specific Dof transcription factors VASCULAR-RELATED DOF1 and VASCULAR-RELATED DOF2 regulate vascular cell differentiation and lignin biosynthesis in Arabidopsis. Plant molecular biology 104, 263–281 (2020).

46 Kubo, H., Kishi, M. & Goto, K. Expression analysis of ANTHOCYANINLESS2 gene in Arabidopsis. Plant science 175, 853–857 (2008).

47 Amanda, D. et al. DEFECTIVE KERNEL1 (DEK1) regulates cell walls in the leaf epidermis. Plant Physiology 172, 2204–2218 (2016).

48 Ingram, P., Dettmer, J., Helariutta, Y. & Malamy, J. E. Arabidopsis Lateral Root Development 3 is essential for early phloem development and function, and hence for normal root system development. The Plant Journal 68, 455–467 (2011).

49 Chen, X. et al. SQUAMOSA promoter-binding protein-like transcription factors: Star players for plant growth and development. Journal of integrative plant biology 52, 946–951 (2010).

50 Denyer, T. et al. Spatiotemporal developmental trajectories in the Arabidopsis root revealed using high-throughput single-cell RNA sequencing. Developmental cell 48, 840–852. e845 (2019).

51 Fang, J. et al. The URL1–ROC5–TPL2 transcriptional repressor complex represses the ACL1 gene to modulate leaf rolling in rice. Plant Physiology 185, 1722–1744 (2021).

52 Horstman, A. et al. AIL and HDG proteins act antagonistically to control cell proliferation. Development 142, 454–464 (2015).

53 Rombolá-Caldentey, B., Rueda-Romero, P., Iglesias-Fernández, R., Carbonero, P. & Oñate-Sánchez, L. Arabidopsis DELLA and two HD-ZIP transcription factors regulate GA signaling in the epidermis through the L1 box cis-element. The Plant Cell 26, 2905–2919 (2014).

54 Yu, L. H. et al. Arabidopsis EDT 1/HDG 11 improves drought and salt tolerance in cotton and poplar and increases cotton yield in the field. Plant biotechnology journal 14, 72–84 (2016).

55 Hong, S.-Y., Kim, O.-K., Kim, S.-G., Yang, M.-S. & Park, C.-M. Nuclear import and DNA binding of the ZHD5 transcription factor is modulated by a competitive peptide inhibitor in Arabidopsis. Journal of Biological Chemistry 286, 1659–1668 (2011).

56 Rosado, D., Ackermann, A., Spassibojko, O., Rossi, M. & Pedmale, U. V. WRKY transcription factors and ethylene signaling modify root growth during the shade-avoidance response. Plant Physiology 188, 1294–1311 (2022).

57 Brockington, S. F. et al. Evolutionary analysis of the MIXTA gene family highlights potential targets for the study of cellular differentiation. Molecular Biology and Evolution 30, 526–540 (2013).

58 Wolfe, K. H., Gouy, M., Yang, Y.-W., Sharp, P. M. & Li, W.-H. Date of the monocot-dicot divergence estimated from chloroplast DNA sequence data. Proceedings of the National Academy of Sciences 86, 6201–6205 (1989).

59 Woolfe, A. et al. Highly conserved non-coding sequences are associated with vertebrate development. PLoS biology 3, e7 (2005).

60 Babarinde, I. A. & Saitou, N. Genomic locations of conserved noncoding sequences and their proximal protein-coding genes in mammalian expression dynamics. Molecular biology and evolution 33, 1807–1817 (2016).

61 Song, B. et al. Conserved noncoding sequences provide insights into regulatory sequence and loss of gene expression in maize. Genome research 31, 1245–1257 (2021).

62 Nelson, A. C. & Wardle, F. C. Conserved non-coding elements and cis regulation: actions speak louder than words. Development 140, 1385–1395 (2013).

63 Hendelman, A. et al. Conserved pleiotropy of an ancient plant homeobox gene uncovered by cis-regulatory dissection. Cell 184, 1724–1739. e1716 (2021).

64 Pennacchio, L. A. et al. In vivo enhancer analysis of human conserved non-coding sequences. Nature 444, 499–502 (2006).

65 McEwen, G. K. et al. Ancient duplicated conserved noncoding elements in vertebrates: a genomic and functional analysis. Genome research 16, 451–465 (2006).

66 Turner, E. E. & Cox, T. C. Genetic evidence for conserved non-coding element function across species–the ears have it. Frontiers in physiology 5, 7 (2014).

67 Nolte, M. J. et al. Functional analysis of limb transcriptional enhancers in the mouse. Evolution & development 16, 207–223 (2014).

68 Salvi, S. et al. Conserved noncoding genomic sequences associated with a flowering-time quantitative trait locus in maize. Proceedings of the National Academy of Sciences 104, 11376–11381 (2007).

69 Vierstra, J. et al. Mouse regulatory DNA landscapes reveal global principles of cis-regulatory evolution. Science 346, 1007–1012 (2014).

70 Leypold, N. A. & Speicher, M. R. Evolutionary conservation in noncoding genomic regions. Trends in Genetics 37, 903–918 (2021).

71 Ciren, D., Zebell, S. & Lippman, Z. B. Extreme restructuring of cis-regulatory regions controlling a deeply conserved plant stem cell regulator. PLoS Genetics 20, e1011174 (2024).

72 Liu, L. et al. Enhancing grain-yield-related traits by CRISPR–Cas9 promoter editing of maize CLE genes. Nature Plants 7, 287–294 (2021).

73 Cao, R. et al. Role of histone H3 lysine 27 methylation in Polycomb-group silencing. Science 298, 1039–1043 (2002).

74 Xiao, J. et al. Cis and trans determinants of epigenetic silencing by Polycomb repressive complex 2 in Arabidopsis. Nature genetics 49, 1546–1552 (2017).

75 Schmitz, R. J., Grotewold, E. & Stam, M. Cis-regulatory sequences in plants: Their importance, discovery, and future challenges. The plant cell 34, 718–741 (2022).

76 76 Ouyang, W., et al. Haplotype mapping of H3K27me3-associated chromatin interactions defines topological regulation of gene silencing in rice. Cell Reports 42 (2023).

77 Minnoye, L. et al. Chromatin accessibility profiling methods. Nature Reviews Methods Primers 1, 10 (2021).

78 Zhang, T.-Q., Chen, Y., Liu, Y., Lin, W.-H. & Wang, J.-W. Single-cell transcriptome atlas and chromatin accessibility landscape reveal differentiation trajectories in the rice root. Nature communications 12, 2053 (2021).

79 Yu, Y., Zhang, H., Long, Y., Shu, Y. & Zhai, J. Plant public RNA-seq database: a comprehensive online database for expression analysis of∼ 45 000 plant public RNA-seq libraries. Plant Biotechnology Journal 20, 806 (2022).

80 Bai, X. et al. Duplication of an upstream silencer of FZP increases grain yield in rice. Nature Plants 3, 885–893 (2017).

81 Wang, Z. et al. AraENCODE: A comprehensive epigenomic database of Arabidopsis thaliana. Molecular Plant 16, 1113–1116 (2023).

82 Tonosaki, K. & Kinoshita, T. Possible roles for polycomb repressive complex 2 in cereal endosperm. Frontiers in plant science 6, 144 (2015).

83 Tan, F.-Q. et al. A coiled-coil protein associates Polycomb Repressive Complex 2 with KNOX/BELL transcription factors to maintain silencing of cell differentiation-promoting genes in the shoot apex. The Plant Cell 34, 2969–2988 (2022).

84 Zhao, L. et al. Integrative analysis of reference epigenomes in 20 rice varieties. Nature communications 11, 2658 (2020).

85 Zhang, J. et al. Building two indica rice reference genomes with PacBio long-read and Illumina paired-end sequencing data. Scientific data 3, 1–8 (2016).

86 Huang, Y. et al. OsNCED5, a 9-cis-epoxycarotenoid dioxygenase gene, regulates salt and water stress tolerance and leaf senescence in rice. Plant science 287, 110188 (2019).

87 Goel, M. et al. The vast majority of somatic mutations in plants are layer-specific. Genome Biology 25, 1–18 (2024).

88 Andrews, G. et al. Mammalian evolution of human cis-regulatory elements and transcription factor binding sites. Science 380, eabn7930 (2023).

89 Engelhorn, J. et al. Phenotypic variation in maize can be largely explained by genetic variation at transcription factor binding sites. bioRxiv, 2023.2008. 2008.551183 (2023).

90 Zhao, T. & Schranz, M. E. Network-based microsynteny analysis identifies major differences and genomic outliers in mammalian and angiosperm genomes. Proceedings of the National Academy of Sciences 116, 2165–2174 (2019).

91 Villar, D. et al. Enhancer evolution across 20 mammalian species. Cell 160, 554–566 (2015).

92 Reineke, A. R., Bornberg-Bauer, E. & Gu, J. Evolutionary divergence and limits of conserved non-coding sequence detection in plant genomes. Nucleic acids research 39, 6029–6043 (2011).

93 Kaplinsky, N. J., Braun, D. M., Penterman, J., Goff, S. A. & Freeling, M. Utility and distribution of conserved noncoding sequences in the grasses. Proceedings of the National Academy of Sciences 99, 6147–6151 (2002).

94 Guo, H. & Moose, S. P. Conserved noncoding sequences among cultivated cereal genomes identify candidate regulatory sequence elements and patterns of promoter evolution. The Plant Cell 15, 1143–1158 (2003).

95 Inada, D. C. et al. Conserved noncoding sequences in the grasses4. Genome Research 13, 2030–2041 (2003).

96 Clark, J. W. & Donoghue, P. C. Whole-genome duplication and plant macroevolution. Trends in plant science 23, 933–945 (2018).

97 Del Pozo, J. C. & Ramirez-Parra, E. Whole genome duplications in plants: an overview from Arabidopsis. Journal of Experimental Botany 66, 6991–7003 (2015).

98 Jump, A. S., Marchant, R. & Peñuelas, J. Environmental change and the option value of genetic diversity. Trends in plant science 14, 51–58 (2009).

99 Wiles, E. T. & Selker, E. U. H3K27 methylation: a promiscuous repressive chromatin mark. Current opinion in genetics & development 43, 31–37 (2017).

100 Zhang, X., Marand, A. P., Yan, H. & Schmitz, R. J. scifi-ATAC-seq: massive-scale single-cell chromatin accessibility sequencing using combinatorial fluidic indexing. Genome Biology 25, 90 (2024).

101 Zheng, G. X. et al. Massively parallel digital transcriptional profiling of single cells. Nature communications 8, 14049 (2017).

102 Ouyang, S. et al. The TIGR rice genome annotation resource: improvements and new features. Nucleic acids research 35, D883–D887 (2007).

103 Satija, R., Farrell, J. A., Gennert, D., Schier, A. F. & Regev, A. Spatial reconstruction of single-cell gene expression data. Nature biotechnology 33, 495–502 (2015).

104 Wolock, S. L., Lopez, R. & Klein, A. M. Scrublet: computational identification of cell doublets in single-cell transcriptomic data. Cell systems 8, 281–291. e289 (2019).

105 Robinson, M. D., McCarthy, D. J. & Smyth, G. K. edgeR: a Bioconductor package for differential expression analysis of digital gene expression data. bioinformatics 26, 139–140 (2010).

106 Crow, M., Paul, A., Ballouz, S., Huang, Z. J. & Gillis, J. Characterizing the replicability of cell types defined by single cell RNA-sequencing data using MetaNeighbor. Nature communications 9, 884 (2018).

107 Rodriques, S. G. et al. Slide-seq: A scalable technology for measuring genome-wide expression at high spatial resolution. Science 363, 1463–1467 (2019).

108 Stickels, R. R. et al. Highly sensitive spatial transcriptomics at near-cellular resolution with Slide-seqV2. Nature biotechnology 39, 313–319 (2021).

109 Satpathy, A. T. et al. Massively parallel single-cell chromatin landscapes of human immune cell development and intratumoral T cell exhaustion. Nature biotechnology 37, 925–936 (2019).

110 Institute, B. Picard toolkit. Broad Institute, GitHub repository (2019).

111 Smit, A. F. Repeat-Masker Open-3.0. http://www.repeatmasker.org (2004).

112 Camacho, C. et al. BLAST+: architecture and applications. BMC bioinformatics 10, 1–9 (2009).

113 Zhang, Y. et al. Model-based analysis of ChIP-Seq (MACS). Genome biology 9, 1–9 (2008).

114 DeBruine, Z. J., Melcher, K. & Triche Jr, T. J. Fast and robust non-negative matrix factorization for single-cell experiments. bioRxiv, 2021.2009. 2001.458620 (2021).

115 McInnes, L., Healy, J. & Melville, J. Umap: Uniform manifold approximation and projection for dimension reduction. arXiv preprint arXiv:1802.03426 (2018).

116 Zong, J. et al. A rice single cell transcriptomic atlas defines the developmental trajectories of rice floret and inflorescence meristems. New Phytologist 234, 494–512 (2022).

117 Hua, L. et al. The bundle sheath of rice is conditioned to play an active role in water transport as well as sulfur assimilation and jasmonic acid synthesis. The Plant Journal 107, 268–286 (2021).

118 Itoh, J.-I. et al. Genome-wide analysis of spatiotemporal gene expression patterns during early embryogenesis in rice. Development 143, 1217–1227 (2016).

119 Wu, T.-Y., Müller, M., Gruissem, W. & Bhullar, N. K. Genome wide analysis of the transcriptional profiles in different regions of the developing rice grains. Rice 13, 1–19 (2020).

120 Van Dijk, D. et al. Recovering gene interactions from single-cell data using data diffusion. Cell 174, 716–729. e727 (2018).

121 Korsunsky, I. et al. Fast, sensitive and accurate integration of single-cell data with Harmony. Nature methods 16, 1289–1296 (2019).

122 Pettkó-Szandtner, A. et al. Core cell cycle regulatory genes in rice and their expression profiles across the growth zone of the leaf. Journal of plant research 128, 953–974 (2015).

123 Schep, A. N. et al. Structured nucleosome fingerprints enable high-resolution mapping of chromatin architecture within regulatory regions. Genome research 25, 1757–1770 (2015).

124 Pattengale, N. D., Alipour, M., Bininda-Emonds, O. R., Moret, B. M. & Stamatakis, A. in Research in Computational Molecular Biology: 13th Annual International Conference, RECOMB 2009, Tucson, AZ, USA, May 18-21, 2009. Proceedings 13. 184-200 (Springer).

125 Bolger, A. M., Lohse, M. & Usadel, B. Trimmomatic: a flexible trimmer for Illumina sequence data. Bioinformatics 30, 2114–2120 (2014).

126 Servant, N. et al. HiC-Pro: an optimized and flexible pipeline for Hi-C data processing. Genome biology 16, 1–11 (2015).

127 Kaul, A., Bhattacharyya, S. & Ay, F. Identifying statistically significant chromatin contacts from Hi-C data with FitHiC2. Nature protocols 15, 991–1012 (2020).

128 Quinlan, A. R. & Hall, I. M. BEDTools: a flexible suite of utilities for comparing genomic features. Bioinformatics 26, 841–842 (2010).

129 Schep, A. N., Wu, B., Buenrostro, J. D. & Greenleaf, W. J. chromVAR: inferring transcription-factor-associated accessibility from single-cell epigenomic data. Nature methods 14, 975–978 (2017).

130 Dijk, D. v., et al. MAGIC: A diffusion-based imputation method reveals gene-gene interactions in single-cell RNA-sequencing data. BioRxiv, 111591 (2017).

131 Alpert, A., Moore, L. S., Dubovik, T. & Shen-Orr, S. S. Alignment of single-cell trajectories to compare cellular expression dynamics. Nature methods 15, 267–270 (2018).

132 Jin, J. et al. PlantTFDB 4.0: toward a central hub for transcription factors and regulatory interactions in plants. Nucleic acids research, gkw982 (2016).

133 Grant, C. E., Bailey, T. L. & Noble, W. S. FIMO: scanning for occurrences of a given motif. Bioinformatics 27, 1017–1018 (2011).

134 Castro-Mondragon, J. A. et al. JASPAR 2022: the 9th release of the open-access database of transcription factor binding profiles. Nucleic acids research 50, D165–D173 (2022).

135 Wagner-Menghin, M. M. Binomial test. Encyclopedia of statistics in behavioral science (2005).

136 Bolstad, B. M. & Bolstad, M. B. M. Package ‘preprocessCore’. (2013).

137 Lovell, J. T. et al. GENESPACE tracks regions of interest and gene copy number variation across multiple genomes. Elife 11, e78526 (2022).

138 Sayers, E. W. et al. Database resources of the national center for biotechnology information. Nucleic acids research 49, D10 (2021).

139 Shen, W., Le, S., Li, Y. & Hu, F. SeqKit: a cross-platform and ultrafast toolkit for FASTA/Q file manipulation. PloS one 11, e0163962 (2016).

140 Pollard, K. S., Hubisz, M. J., Rosenbloom, K. R. & Siepel, A. Detection of nonneutral substitution rates on mammalian phylogenies. Genome research 20, 110–121 (2010).

141 Siepel, A. & Haussler, D. Phylogenetic estimation of context-dependent substitution rates by maximum likelihood. Molecular biology and evolution 21, 468–488 (2004).

142 Langmead, B. & Salzberg, S. L. Fast gapped-read alignment with Bowtie 2. Nature methods 9, 357–359 (2012).

143 Li, H. et al. The sequence alignment/map format and SAMtools. bioinformatics 25, 2078–2079 (2009).

144 Stovner, E. B. & Sætrom, P. epic2 efficiently finds diffuse domains in ChIP-seq data. Bioinformatics 35, 4392–4393 (2019).

145 Xie, L. et al. RiceENCODE: A comprehensive epigenomic database as a rice Encyclopedia of DNA Elements. Molecular Plant 14, 1604–1606 (2021).

146 Chen, S., Zhou, Y., Chen, Y. & Gu, J. fastp: an ultra-fast all-in-one FASTQ preprocessor. Bioinformatics 34, i884–i890 (2018).

147 Li, H. & Durbin, R. Fast and accurate short read alignment with Burrows–Wheeler transform. bioinformatics 25, 1754–1760 (2009).

148 McKenna, A. et al. The Genome Analysis Toolkit: a MapReduce framework for analyzing next-generation DNA sequencing data. Genome research 20, 1297–1303 (2010).

149 Tian, T. et al. agriGO v2. 0: a GO analysis toolkit for the agricultural community, 2017 update. Nucleic acids research 45, W122-W129 (2017).

150 Hofmeister, B. T. & Schmitz, R. J. Enhanced JBrowse plugins for epigenomics data visualization. BMC bioinformatics 19, 1–6 (2018).

